# The complexity and dynamics of *in organello* translation assessed by high-resolution mitochondrial ribosome profiling

**DOI:** 10.1101/2023.07.19.549812

**Authors:** Taisei Wakigawa, Mari Mito, Haruna Yamashiro, Kotaro Tomuro, Haruna Tani, Kazuhito Tomizawa, Takeshi Chujo, Asuteka Nagao, Takeo Suzuki, Fan-Yan Wei, Yuichi Shichino, Tsutomu Suzuki, Shintaro Iwasaki

**Affiliations:** RNA Systems Biochemistry Laboratory, RIKEN Cluster for Pioneering Research, Wako, Saitama 351-0198, Japan; Department of Computational Biology and Medical Sciences, Graduate School of Frontier Sciences, The University of Tokyo, Kashiwa, Chiba 277-8561, Japan; Department of Modomics Biology and Medicine, Institute of Development, Aging and Cancer, Tohoku University, Sendai, Miyagi 980-8575, Japan; Department of Molecular Physiology, Faculty of Life Sciences, Kumamoto University, Kumamoto, Kumamoto 860-8556, Japan; Department of Chemistry and Biotechnology, Graduate School of Engineering, The University of Tokyo, Bunkyo-ku, Tokyo, 113-8656, Japan; Department of Medical Biochemistry, Graduate School of Medicine, University of the Ryukyus, Nishihara, Okinawa 903-0215, Japan

**Author notes:** Correspondence should be addressed to S.I. These authors contributed equally (Taisei Wakigawa, Mari Mito).

## Abstract

Since mitochondrial translation serves the essential subunits of the OXPHOS complex that produces ATP, exhaustive, quantitative, and high-resolution delineation of mitoribosome traversal is needed. Here, we developed a technique for high-resolution mitochondrial ribosome profiling and revealed the intricate regulation of mammals *in organello* translation. Our approach assessed the stoichiometry and kinetics of mitochondrial translation flux, such as the number of mitoribosomes on a transcript and the elongation rate, initiation rate, and lifetime rounds of translation of individual transcripts. We also surveyed the impacts of modifications at the anticodon stem loop in mt-tRNAs, including all possible modifications at the 34th position, by deleting the corresponding enzymes and harnessing patient-derived cells. Moreover, a retapamulin-assisted derivative and mito-disome profiling revealed cryptic translation initiation sites at subcognate codons and programmed mitoribosome collision sites across the mitochondrial transcriptome. Our work provides a useful platform for investigating protein synthesis within the energy powerhouse of the cell.

## Introduction

In eukaryotic cells, mitochondrial symbiogenesis provides another protein synthesis system in the organelle ^1^, in addition to one in the cytosol ^2,3^. Although the human mitochondrial genome possesses only 13 ORFs, translation in the mitochondrial matrix is indispensable for cells since these mRNAs encode a subunit of oxidative phosphorylation (OXPHOS) that is synthesize ATP ^4^. The importance of mitochondrial translation integrity has been illustrated by the pathological outcome of its dysregulation ^5–7^.

A variety of techniques have been developed to investigate translation in mitochondria ^8^. Among them, mitochondria-optimized ribosome profiling (or Ribo-Seq), which is based on deep sequencing of ribosome-protected RNA fragments remaining after RNase digestion ^9^, has been a powerful tool (MitoRibo-Seq) ^10–16^. Technically, MitoRibo-Seq requires the depletion of cytosolic ribosome (cytoribosome) footprints; otherwise, they occupy large sequencing space. For this purpose, a sucrose density gradient with ultracentrifugation has been applied to enrich the mitoribosome-footprint complex ^10–16^. A drawback of this strategy is its limitation in handling many samples (*i.e.*, the maximum number of samples is limited to the 6 tubes that the ultracentrifuge rotor holds) as well as its time-consuming nature (∼1 d). In yeast, another approach involves epitope tagging of mitoribosomal proteins and further immunopurification of mitoribosomes. Although this technique should be versatile in mammals, the development of cell lines or animals expressing tagged mitoribosomes is a technical hurdle. Thus, there is still much room for improvement in MitoRibo-Seq as a simple, low input-tailored, and sample number-scalable tool.

Related to the issues described above, our knowledge of mitochondrial translation kinetics in cells has been limited. This is in stark contrast to the case for cytosolic translation, where kinetic information has been generated by *in vivo* single-molecule imaging ^17–23^ and cytosolic ribosome profiling ^24–26^.

To expand our understanding of mitochondrial translation, we developed MitoIP-Thor-Ribo-Seq. Mitochondrial immunoprecipitation (MitoIP) with an antibody targeting the outer membrane protein TOM22 allows rapid purification of mitochondrial ribosomes with parallel handling of multiple samples simultaneously. Moreover, the implementation of an RNA-dependent RNA amplification technique ^27^ has enabled library preparation from low-input materials after MitoIP. By combining a retapamulin-mediated mitochondrial ribosome (mitoribosome) run-off assay and a calibration technique ^26^, we measured key kinetic and stoichiometric parameters of *in organello* translation, including translation initiation and elongation rates. Extremely slow elongation and recycling events were further revealed by sequencing footprints generated by colliding mitochondrial disomes (mito-disomes). Moreover, retapamulin-assisted MitoIP-Thor-Ribo-Seq revealed subcognate cryptic translation initiation sites in the mitochondrial transcriptome. We further employed our method to study the role of mitochondrial tRNA (mt-tRNA) modifications at the 34th and 37th positions. This method provides a versatile and high-resolution tool for monitoring *in organello* protein synthesis, serving a valuable data resource for future studies.

## Results

### Development of MitoIP-Thor-Ribo-Seq

To overcome the issues of mitochondrial ribosome profiling mentioned above, we employed mitochondrial immunoprecipitation (MitoIP) with an anti-TOM22 antibody and then treated the mitoribosome-enriched material with RNase to generate the footprints (Figure 1A). The MitoIP approach effectively concentrated mitoribosomes (55S, 39S, and 28S) from HEK293 cells, as detected in the sucrose density gradient, whereas the signal of mitochondrial ribosome complexes could not be detected in total cell lysates due to the large excess of cytosolic ribosomes (Figure 1B).

**Figure 1.**
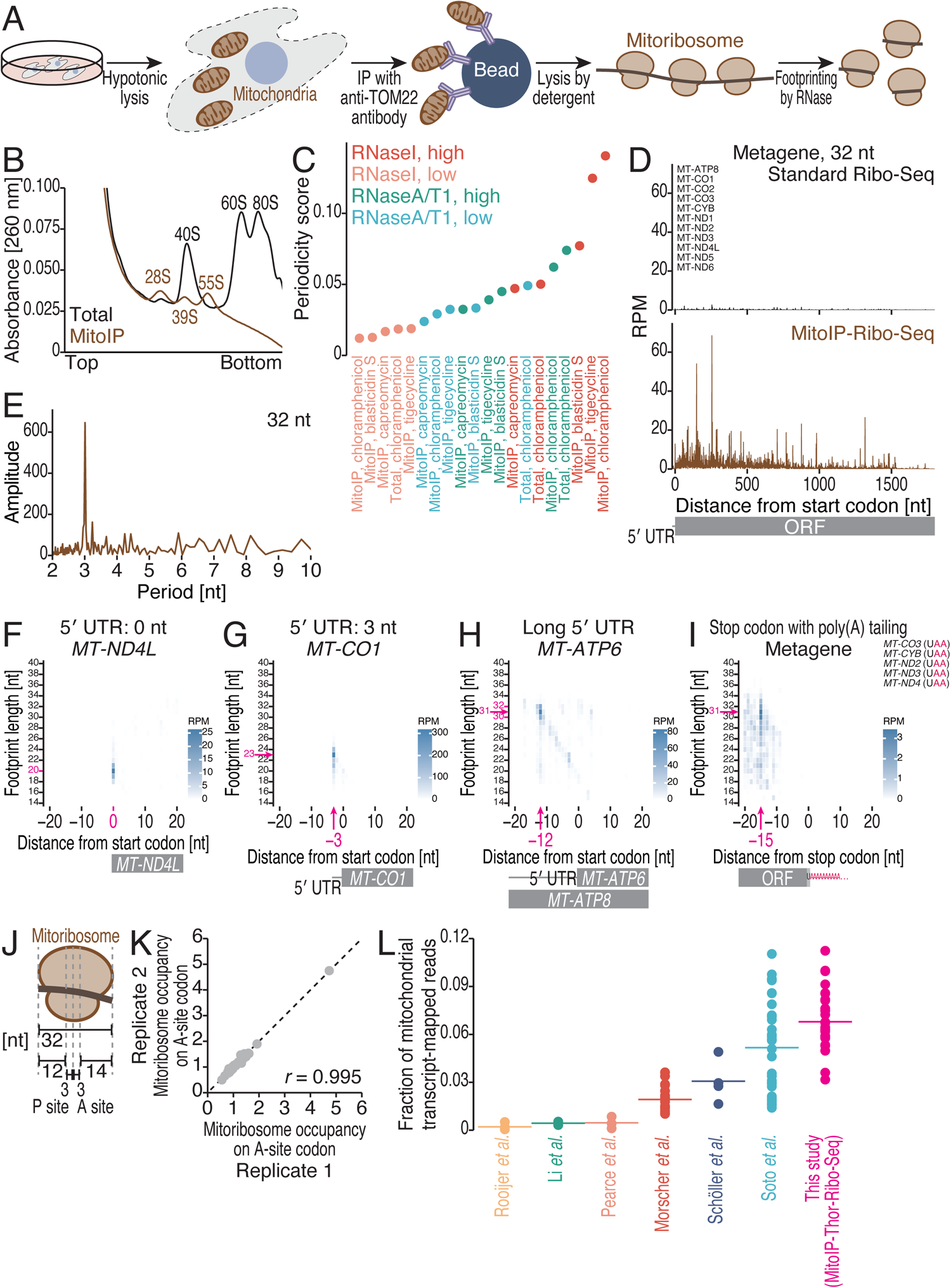
Mitochondrial translation-tailored ribosome profiling with sample number scalability and low-input compatibility. (A) Schematic of mitochondrial immunoprecipitation and subsequent ribosome profiling (MitoIP-Ribo-Seq). (B) Sucrose density gradient ultracentrifugation for the separation of cytosolic and mitochondrial ribosome complexes. (C) Periodicity scores across the indicated conditions of MitoIP-Ribo-Seq. The periodicity score was defined as the sum of the information content for the 3-nt periodicity at each read length, weighted by the fraction of the read length in the library. Figure S1F-G shows schematic representations of the information content for the 3-nt periodicity and the periodicity score calculation scheme. (D) Metagene plots for the 5′ ends of 32-nt mitoribosome footprints around start codons (the first nucleotide of the start codon was set to 0) in MitoIP-Ribo-Seq (brown) and standard Ribo-Seq (black). *MT-ATP6* and *MT-ND4* were excluded from the analysis because their ORF start codons overlapped with those of *MT-ATP8* and *MT-ND4L*, respectively. Notably, due to short footprints at the start codon and downstream codons (see F and G), the read coverage in such a region was underrepresented. RPM, reads per million mapped reads. (E) Discrete Fourier transform of mitoribosome footprints mapped to the first 300-nt region from the start codon of the indicated ORFs. (F-H) Plots of the 5′ ends of the mitoribosome footprint along the length around the start codon (the first nucleotide of the start codon was set to 0) for the indicated transcripts. The color scale indicates read abundance. (I) Metagene plots for the 5′ ends of mitoribosome footprints along the length around stop codons (the first nucleotide of the stop codon was set to 0) for the transcripts whose stop codons were generated by poly(A) tailing. The color scale indicates the read abundance. (J) Schematic representation of the relative position of the P site and A site on the mitoribosome footprints. (K) Scatter plot of the mitoribosome occupancy at all the codons across mitochondrial ORFs in replicates. *r*, Pearson’s correlation coefficient. (L) Comparison of the fraction of mitochondrial transcript-mapped reads between our methods and published data. See also Figure S1 and Table S1.

We systematically optimized the library construction procedures for mitoribosome footprints (or MitoIP-Ribo-Seq) using HEK293 cell lysates. Since ribosome isolation after RNase treatment was crucial for enriching footprints and excluding non-footprinting RNA fragments, we compared a sucrose cushion with ultracentrifugation and a size-exclusion gel-filtration spin column ^28^ and found that the former was more effective (Figure S1A-B, represented by the depletion of TOM20 and GAPDH). Given that RNase selection and treatment conditions define the quality of ribosome profiling data ^29–32^, we optimized the application in terms of RNase type (RNase I or a mixture of RNase A and RNase T1) and concentration (low or high). Moreover, we tested ribosome inhibitors in MitoIP-Ribo-Seq (Figure S1C-D), since compounds that halt ribosome movement along mRNAs are also pivotal factors in these experiments ^33–39^. Notably, we omitted the drug treatment of the cell culture and instead added the drugs to the buffer for cell lysis (postlysis treatment) ^40^ to avoid potential artifacts introduced by pretreatment in cell culture ^35–37,39^. In addition, we harnessed the recently developed Ribo-FilterOut technique to reduce rRNA contamination and increase the sequencing space (Figure S1E) ^26^. To evaluate the quality of the ribosome profiling data, we defined the periodicity score (Figure S1F-G) to evaluate the 3-nt periodicity, which is a hallmark of ribosome profiling data with high codon resolution (Figure S1H).

Ultimately, we found that MitoIP-Ribo-Seq with chloramphenicol-mediated mitoribosome elongation block and a high concentration of RNase I had the greatest results (Figures 1C and S1I), providing mitoribosome footprints that peaked at 31-32 nt (Figure S1H). Compared to standard Ribo-Seq, optimized MitoIP-Ribo-Seq provided a significant enrichment of mitoribosome footprints along ORFs (Figure 1D) with fine triplet periodicity (Figure 1E).

The mitoribosome footprint in our datasets reflected the characteristic features of mitochondrial mRNAs. As 8 of 13 mRNAs do not have a 5′ UTR ^41^, 55S mitoribosomes assembled on the start codons at the very 5′ ends of mRNAs should not have the RNA region to be protected by the mitoribosomes ^16^. Consistent with this scenario, we observed shorter 20-nt footprints at the 5′ ends of mRNAs without 5′ UTRs (Figures 1F and S1J-K). We found that the footprints at the start codons with longer 5′ UTRs were extended; for example, we observed 23-nt footprints for *MT-CO1* mRNA, which has a 3-nt 5′ UTR (Figure 1G), and 31-nt and 32-nt footprints for *MT-ATP6* (Figure 1H) and *MT-ND4* (Figure S1L), which have long 5′ UTRs.

Seven of the 13 ORFs require poly(A) tails to generate a complete stop codon sequence (UAA) ^41^. Our MitoIP-Ribo-Seq detected footprints on poly(A)-generated stop codons (Figures 1I and S1M), suggesting that poly(A) tailing is highly efficient and that mitoribosomes predominantly translate mRNAs after polyadenylation.

Given the footprint length and start codon locations that should be in the P site of mitoribosomes (Figures 1F-H and S1J-L), the P-site position in the footprints was determined to be 12 nt downstream from the 5′ ends (Figure 1J). Consistently, we observed the 5′ ends of footprints 15 nt upstream of the stop codons, which are positioned at the A site (Figures 1I and S1M). Taking the 3-nt periodicity (Figure 1E) into account, we concluded that our MitoIP-Ribo-Seq can assess mitoribosome locations at single-codon resolution.

Due to the process of mitochondria isolation, only a small amount of material is ultimately acquired, hampering robust library construction. For example, we could recover materials that yields only 1-2 µg of RNA after MitoIP from ∼10-20 × 10^6^ cells on 15-cm dishes, although stable library preparation requires sample corresponding to 10 µg of RNA ^27^. Thus, we implemented the T7 high-resolution original RNA (Thor)-Ribo-Seq approach, which is based on RNA-dependent RNA amplification ^27^. The application of this technique provided highly reproducible results (Figure 1K and Table S1, Pearson’s correlation of 0.983 on average in replicates), with codon-wise measurements of ribosome occupancy comparable to those by the standard method (Figure S1N). Therefore, we harnessed MitoIP-Thor-Ribo-Seq for downstream analysis. We note that although the chloramphenicol used in our method has been proposed to cause context-selective inhibition of elongation (preference for Ala/Ser and disfavor for Gly) at the penultimate amino acid (or E site) in bacteria ^42–46^, our mitochondrial data did not show such bias (Figure S1N).

Ultimately, compared to published conventional MitoRibo-Seq ^10,11,13–16^, our MitoIP-Thor-Ribo-Seq provided comparable or even superior results in terms of mitoribosome footprint recovery (Figure 1L). Notably, the convenience of parallel mitochondrial isolation for multiple samples and the low input-tailored method allowed us to investigate mitochondrial translation in more than 70 samples in this study.

### Kinetics of mitochondrial translation

During our survey of ribosome inhibitors for mitochondrial translation based on on-gel mitochondrial-specific fluorescent noncanonical amino acid tagging (on-gel mito-FUNCAT) ^47–50^ (Figure S1C-D), we noticed that retapamulin, a compound that traps bacterial ribosomes at the start codon but does not actively elongating ribosomes ^51^, effectively blocked mitochondrial translation. This led us to perform a mitoribosome run-off assay ^24,25^ to measure the mitochondrial translation elongation rate (Figure 2A) — pulse treatment with retapamulin to inhibit translation initiation, the subsequent chase of preloaded mitoribosomes over time, and elongation arrest with chloramphenicol. This analysis should generate mitoribosome-free regions downstream of start codons (Figure 2A). Indeed, we observed a chase time-dependent extension of the region by MitoIP-Thor-Ribo-Seq in HEK293 cells (Figures 2B and S2A). Regression fitting revealed a global mitoribosome elongation rate of 0.515 codon/s (Figure 2C).

**Figure 2.**
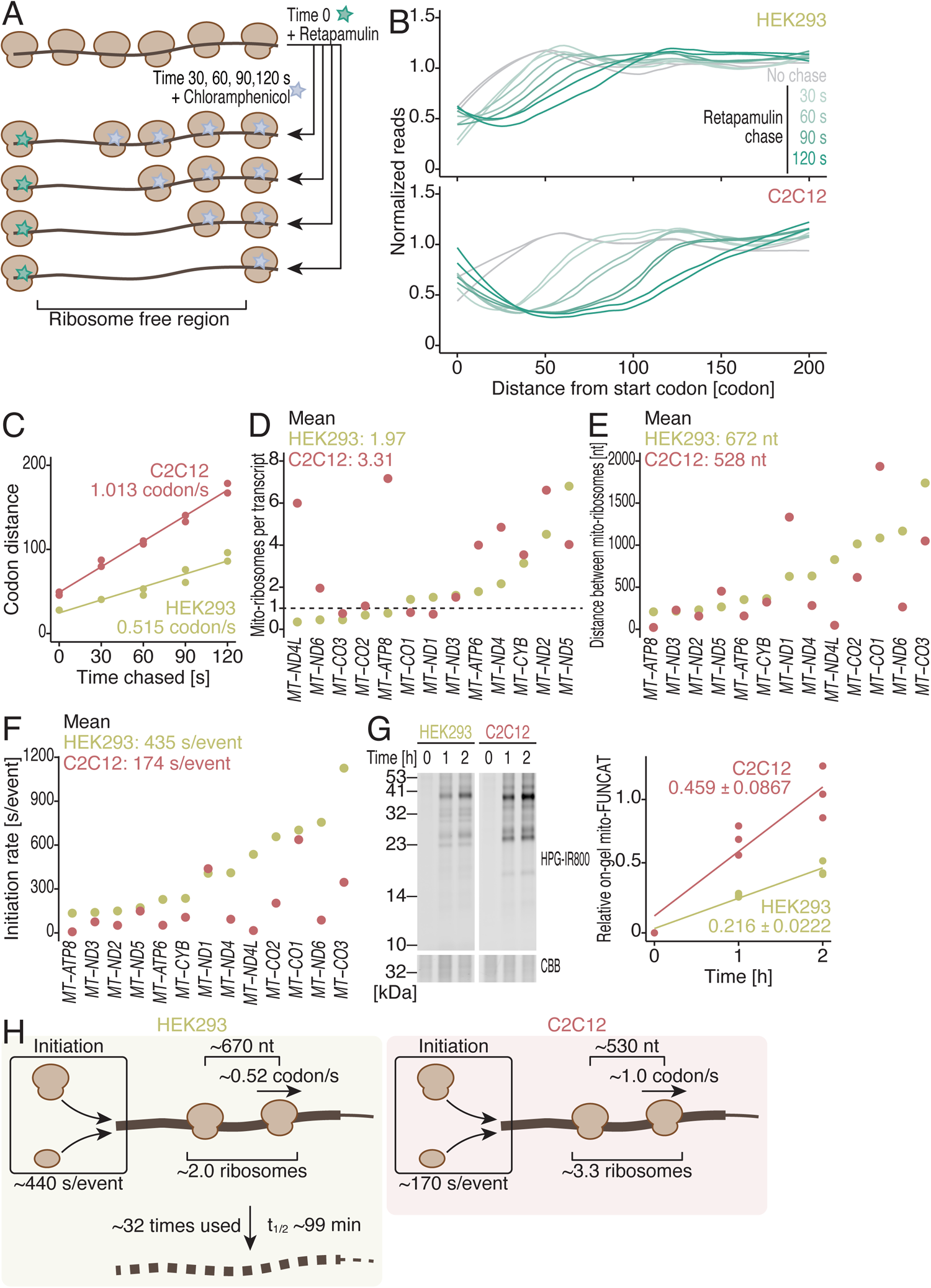
Mitochondrial translation kinetics. (A) Schematic of the mitoribosome run-off assay with retapamulin chase. After a short incubation with retapamulin, the mitoribosomes preloaded on the transcripts were fixed with chloramphenicol. (B) Smoothed metagene plots with footprints around start codons (the first nucleotide of the start codon was set to 0) in the mitoribosome run-off assay for the indicated cell lines. *MT-ATP6* and *MT-ND4* were excluded from the analysis since their ORF start codons overlapped with those of *MT-ATP8* and *MT-ND4L*, respectively. (C) The expansion of the mitoribosome-free area over time by retapamulin chase in the indicated cell lines. Linear regression was used to determine the global elongation rate of mitochondrial translation. (D) The number of mitoribosomes on the indicated ORFs in the indicated cell lines, determined by Ribo-Calibration. (E) The distance between mitoribosomes on the indicated ORFs in the indicated cell lines, determined by Ribo-Calibration. (F) The translation initiation rate of the indicated ORFs in the indicated cell lines. (G) On-gel mito-FUNCAT [labeled with L-homopropargyl glycine (HPG) conjugated with infrared 800 (IR800)] was used to monitor mitochondrial translation over time in the indicated cell lines. The data reported in an earlier study ^50^ were reanalyzed. Coomassie Brilliant Blue (CBB) staining of the total protein was used for the loading control. On the right panel, individual replicates (n = 3, points) are shown. Linear regression was used to determine the mitochondrial protein synthesis rate. Errors represent s.d.s. (H) Schematic summary of the mitochondrial translation kinetics parameters in two cell lines. See also Figure S2 and Table S1-2.

Our recent method for calibrating Ribo-Seq data (Ribo-Calibration) converts ribosome footprint counts to the number of ribosomes on transcripts ^26^. This analysis in HEK293 cells estimated differential mitoribosome numbers on transcripts (Figure 2D), while the number was generally low; on average, ∼2 mitoribosomes were loaded on each ORF. The sparseness was exemplified by the long distances between mitoribosomes (672 nt on average) (Figure 2E). Considering the number of mitoribosomes per transcript, the translation elongation rate calculated above, and the ORF length, we calculated the translation initiation rate for each ORFs (see Materials and Methods for details) ^26^. Generally, mitochondrial translation initiation was extremely slow, at 435 s/event on average (Figure 2F).

To investigate the variability of the kinetics and stoichiometry in mitochondrial translation, we applied the same mitoribosome run-off assay and Ribo-Calibration with mouse myoblast C2C12 cells (Figure 2B-F). We observed significant differences between the two cell lines: a faster elongation rate (1.01 codon/s) and greater translation initiation rate (174 s/event on average). Moreover, mRNA-wise variance in the number of mitoribosomes and in initiation speed was observed (Figure 2D-F).

To validate the translation initiation rates that determine the overall protein synthesis speed, we used mito-FUNCAT ^50^. C2C12 cells were 2.1-fold faster than HEK293 cells (Figure 2G), which is consistent with the results of the translation initiation rates calculated above (∼2.5-fold).

We further measured mRNA half-lives by 5′-bromo-uridine immunoprecipitation chase–deep sequencing (BRIC-Seq) ^26,52,53^ and found a similar range of mRNA half-lives to those previously reported ^54,55^ (Figure S2B), although the numbers for individual mRNAs differed between our study and those studies (Table S2). This measurement allowed us to calculate the lifetime translation cycles before transcript decay; we determined that the average number of mitochondrial mRNAs used for protein synthesis was 31.8 (Figure S2C).

Thus, our analysis revealed the stoichiometric and kinetic parameters of *in organello* translation and their difference in cell types (Figure 2H).

### Widespread cryptic translation initiation sites

Given that retapamulin-mediated mitoribosome arrest occurs at the start codon, we reasoned that long-term incubation with this compound could be harnessed for translation initiation site surveys similar to those conducted for bacterial translation (with retapamulin) ^51^ and cytosolic translation in eukaryotes (with harringtonine or lactimidomycin) ^25,56,57^. As expected, retapamulin-assisted MitoIP-Thor-Ribo-Seq in HEK293 cells revealed sharp read peaks at all the annotated translation initiation sites both at the 5′ ends of mRNAs (Figures 3A-C and S3A-H) and in the middle of transcripts in the polycistronic *MT-ATP8*/*MT-ATP6* (Figure 3A) and *MT-ND4L*/*MT-ND4* mRNAs (Figure 3B). Accordingly, the short footprints that reflect mitoribosomes at the start codons of leaderless mRNAs (Figures 1F-G and S1J-K) were enriched by retapamulin treatment (Figure 3D).

**Figure 3.**
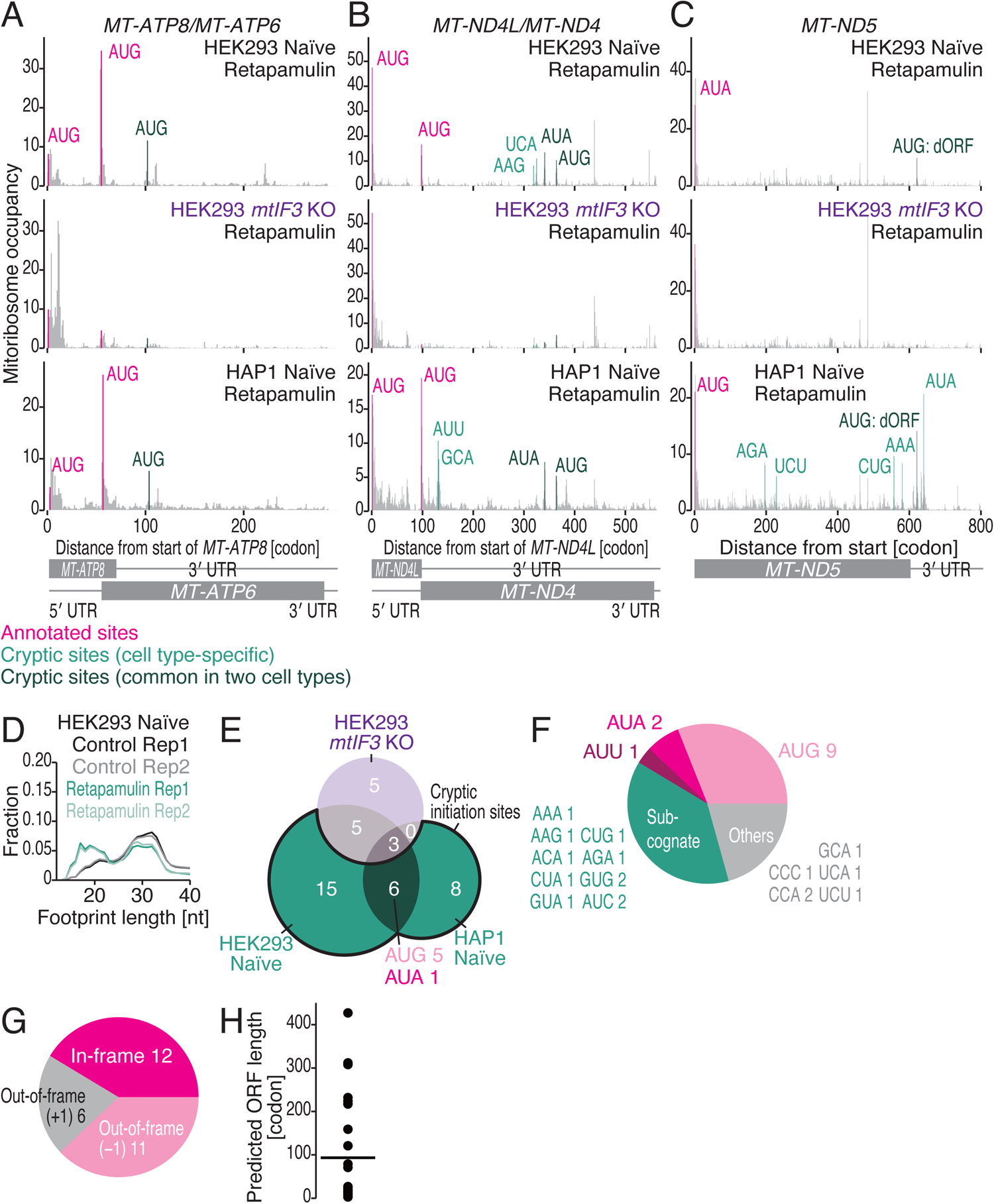
Translation initiation sites survey by retapamulin-assisted MitoIP-Thor-Ribo-Seq. (A-C) Distribution of mitoribosome footprints from retapamulin-treated and control cells along the indicated transcripts. The A-site position of each read is indicated. Codon sequences corresponding to the P site are highlighted. Magenta, annotated initiation sites; peacock green, cryptic initiation sites found in a cell type-specific manner; and dark green, cryptic initiation sites found in both cell lines. See Table S3 for the full list. (D) The fraction of mitoribosome footprints along the read length for the indicated conditions. (E) Venn diagrams for the peak sites found in the indicated cell lines. The subgroup highlighted by the bald black line was defined as cryptic initiation sites. (F-G) Pie charts for codon sequences of the cryptic translation initiation sites (F) and their frame positions relative to the annotated ORFs (G). (H) The length of the ORFs predicted by the cryptic translation initiation sites. See also Figure S3 and Table S1 and S3.

Intriguingly, we also detected the accumulation of mitoribosome footprints at diverse positions in the mitochondrial transcriptome. Thus, we systematically surveyed the cryptic translation initiation sites using retapamulin treatment data. Considering annotated translation initiation sites as true positives, a receiver operating characteristic (ROC) curve defined the threshold of peak calling at 99% specificity (1% false positive rate) (Figure S3I-J) and identified 29 unannotated positions (Figure 3E).

Recent reports have suggested that protein synthesis from polycistronic mRNAs may depend on a unique process for translation initiation ^58,59^; the internal ORFs of *MT-ATP6* and *MT-ND4* exclusively require mitochondrial translation initiation factor (mtIF) 3 but ORFs at the 5′ end of mRNAs does not. Therefore, we reasoned that if the unannotated sites found in the internal positions of mRNAs function as translation initiation sites, their recognition by the mitoribosome is also mediated by mtIF3. For this purpose, we generated a *mtIF3* knockout (KO) cell line (Figure S3K-M) and applied retapamulin-assisted MitoIP-Thor-Ribo-Seq (Figure S3N). Consistent with an earlier report ^59^, mitoribosomes loaded on the start codons in *MT-ATP6* and *MT-ND4* were drastically decreased in *mtIF3* KO, whereas translation initiation on upstream *MT-ATP8* and *MT-ND4L* was not affected (Figure 3A-B). Similar to the start codons of *MT-ATP6* and *MT-ND4*, the majority of the unannotated positions (21 out of 29, 72%) had drastically decreased reads and were no longer called as peak sites (Figure 3E). Here, we excluded the peak sites still found in *mtIF3* KO HEK293 cells for downstream analysis. To further extend our analysis to other cell types, we repeated retapamulin-assisted MitoIP-Thor-Ribo-Seq to HAP1 cells (Figure S3P) and defined 14 sites (Figures 3A-C, E, and S3A-H), taking the overlapping sites found in *mtIF3* KO HEK293 cells into account. Ultimately, our data listed 29 sites in HEK293 and HAP1 cells with only 6 common sites (Figure 3E and Table S3), suggesting a cell type-specific selection of cryptic initiation sites. We noted that these initiation sites were not associated with mitoribosome stalling in general (Figure S3Q-R).

These cryptic translation initiation sites contained AUG, AUA, and AUU codons, as found in annotated start codons, or their subcognate sequences (23 out of 29, 79%) (Figure 3F). Generally, they may synthesize short peptides from the out-of-frame sequences (Figure 3G-H and Table S3), although a subset of in-frame initiation sites may generate N-terminal truncated proteins (Figure 3G-H and Table S3). This set of newly identified translation initiation sites included a recently reported short ORF in the 3′ UTR of *MT-ND5* [*MT-ND5* downstream ORF (*MT-ND5-dORF*)] ^16^ (Figure 3C).

These data revealed the unexpected complexity of start codon recognition in mitochondrial protein synthesis.

### The impacts of modifications in mitochondrial tRNAs on mitoribosome traversal

Harnessing the MitoIP-Thor-Ribo-Seq approach, we assessed the importance of mitochondrial tRNA (mt-tRNA) modifications on mitochondrial translation ^60,61^ (Figure 4A). Here, we focused on all the modifications found at the 34th position [5-formylcytidine (f^5^C), 5-taurinomethyluridine (τm^5^U), 5-taurinomethyl-2-thiouridine (τm^5^s^2^U), and queuosine (Q)] and *N*^6^-threonylcarbamoyladenosine (t^6^A) found at the 37th position (Figures 4A and S4A-D). We used knockout (KO) HEK293T cell lines in which the responsible enzymes were deleted by CRISPR-Cas9, and the mt-tRNA modification status was well characterized by earlier studies (*NSUN3* KO ^62^, *GTPBP3* KO ^63^, *QTRT1* KO ^60^, *QTRT2* KO ^60^, and *OSGEPL1* KO ^64^). In addition, we newly generated *MTU1* KO HEK293 cells (Figure S4E), which ensured the complete loss of τm^5^s^2^U34 (Figure S4F). MitoIP-Thor-Ribo-Seq in these cells (Figure S4G-K) revealed decreased mitoribosome traversal depending on the codon at the A site (Figure 4B) and P site (Figure S4M). The accumulation of mitoribosome occupancy strongly corresponded to the codons that were decoded by the decorated mt-tRNAs (see below). For mRNA-wise analysis, we quantified polarity scores ^65^, which reflect the positional bias of footprints in ORFs (Figure S4N), and the 3-nt periodicity (bit) (Figure S1F). The loss of mt-tRNA modification assessed in this study often led to mitoribosome partitioning toward the 5′ end in some specific ORFs and the associated reduction in the 3-nt periodicity, which may suggest frameshifts and/or premature terminations (Figure 4C-G).

**Figure 4.**
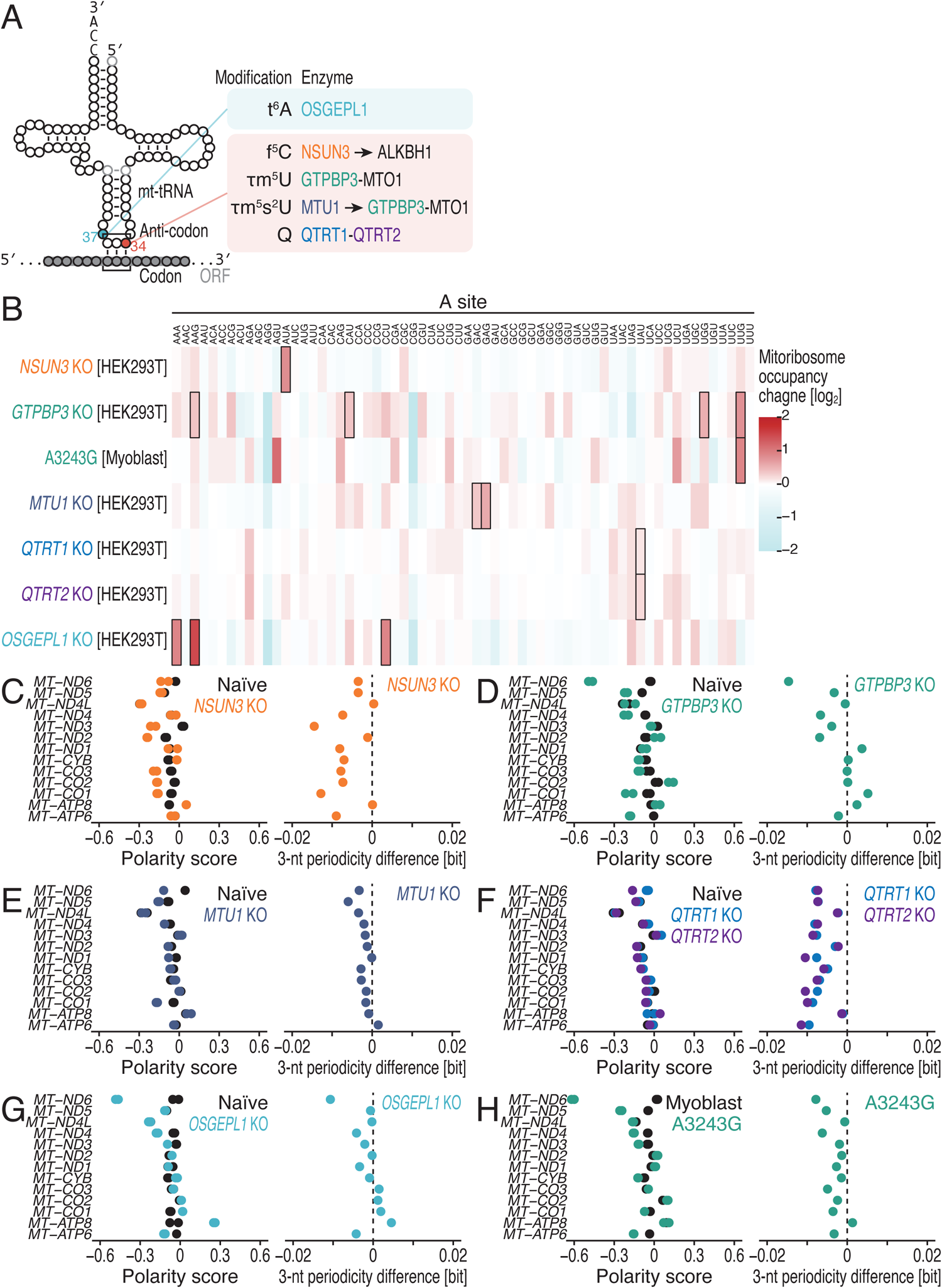
An overview of the translation alterations induced by the loss of the mt-tRNA modifications. (A) Schematic of the modified nucleotides and their positions on mt-tRNAs investigated in this study. Responsible enzymes for the modifications are shown. (B) A heatmap for the mitoribosome occupancy changes at A-site codons in the indicated cell lines. The color scale indicates mitoribosome occupancy change. The remarkable mitoribosome occupancy increases found in the mutant cells are highlighted in black rectangles. (C-H) Polarity scores of the indicated ORFs in the indicated cell lines (left). The information content for the 3-nt periodicity of the reads from the indicated ORFs was calculated and then the changes by the gene knockout were calculated (right). For the polarity scores, the data points from two replicates are shown individually. For the 3-nt periodicity difference, the means of two replicates are shown. See also Figure S4-9 and Table S1.

### f^5^C34 for AUA codons

f^5^C34 is found in mt-tRNA^Met^, which decodes the methionine AUA and AUG codons^1,41,61^. This modification is catalyzed in 3 steps by 2 enzymes (Figures 4A and S4A) ^62,66–68^: cytidine (C) conversion to 5-methylcytidine (m^5^C) (methylation) by NSUN3, followed by m^5^C conversion to 5-hydroxymethylcytidine (hm^5^C) (hydroxylation) and hm^5^C conversion to f^5^C (oxidation) by ALKBH1 ^62,66–68^. Although this modification has been suggested to be essential for AUA recognition by *in vitro* systems ^69,70^, its impact on *in organello* translation remains elusive. Consistent with the proposed functions of the modifications in decoding, MitoIP-Thor-Ribo-Seq revealed a prominent accumulation of mitoribosome footprints on the AUA codon at the A site by *NSUN3* deficiency (Figures 4B and S5A). In stark contrast, alterations in mitoribosome occupancy at the AUG codon were limited (Figures 4B and S5A). These data highlighted the significance of f^5^C34 on mt-tRNA^Met^; the hypomodification led to efficiently decoding AUA at the A site (Figure S9A).

Moreover, our data revealed mitoribosome accumulation on AUA at the P Site (Figure S4M), suggesting that f5C34 plays a role in peptidyl transfer or mitoribosome translocation. Given that the AUA codon (166 sites) is more frequently used than the AUG codon (40 sites) in mitochondrial mRNAs ^62^, two consecutive AUA codons may occur in mitochondrial mRNAs (10 sites). Indeed, the AUA-AUA dicodons in the P and A sites further enhanced overall mitoribosome stalling (Figure 5A). This phenotype was exemplified by remarkable mitoribosome footprint accumulation in an AUA-AUA dicodon in *MT-ND5* (Figure 5B).

**Figure 5.**
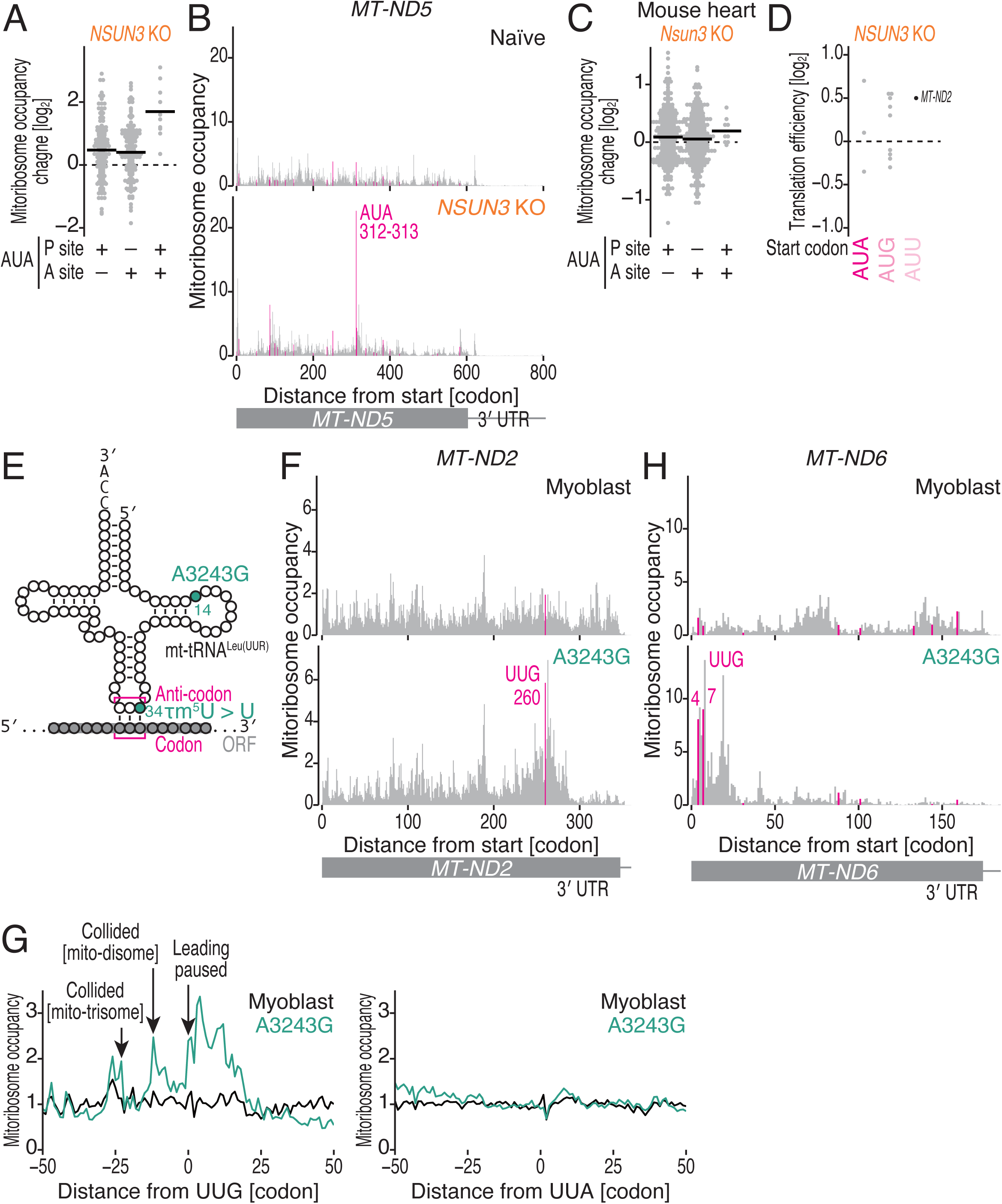
Translational dysregulation by the loss of f^5^C modification and in MELAS patient myoblasts. (A) Mitoribosome occupancy changes at the P-site AUA, A-site AUA, and PA-site AUA-AUA codons in *NSUN3* KO. (B) Distribution of mitoribosome footprints from naïve and *NSUN3* KO cells along the indicated transcripts. The A-site position of each read is indicated. AUA codons are highlighted. (C) Mitoribosome occupancy changes at the P-site AUA, A-site AUA, and PA-site AUA-AUA codons in heart tissues from *Nsun3* KO mice. (D) Translation efficiency changes measured by standard Ribo-Seq and RNA-Seq for mitochondrial transcripts in *NSUN3* KO. Transcripts are sorted according to their start codon sequences. (E) Schematic of the A3243G mutation in mt-tRNA^Leu^ ^(UUR)^. (F and H) Distribution of mitoribosome footprints from naïve and A3243G myoblasts along the indicated transcripts. The A-site position of each read is indicated. UUG codons are highlighted. (G) Metagene plots for mitoribosome footprints from naïve and A3243G myoblasts around UUG codons (left) and UUA codons (right). See also Figures S4-6 and S9 and Table S1.

In addition to the strong pinpoint effect on specific codons, the loss of the f^5^C34 modification may also shift the distribution of the mitoribosome toward the 5′ region (Figure 4C). *MT-CO1*, which has a cluster of AUA codons from positions 50-100, provided such an example (Figure S5B).

To further extend our approach from cell culture to *in vivo* tissue, we employed hearts from *Nsun*3 KO mice ^71^ (Figures 5C and S5C-H). The mitoribosome footprints from the mouse tissue also showed shorter mitoribosome footprints at start codons on mRNAs with absent or short 5′ UTRs (Figure S5C-D). Our mouse heart data recapitulated ribosome stalling at the AUA-AUA dicodons, although the effect was less pronounced than that in cell culture (Figure 5C). We observed that the distribution of the mitoribosome footprint was biased toward the 5′ ends of transcripts possessing AUA clusters at the early ORF, such as those of *MT-CO1* and *MT-ND5* (Figure S5F-H). These data indicated the functional conservation of f^5^C34 in mammals and demonstrated the potential of our approach for *in vivo* study.

In addition to participating in elongation, mt-tRNA^Met^ should be used as the initiator tRNA ^1,61^. To monitor the effect of modifications on translation initiation, we conducted standard Ribo-Seq (without MitoIP) and RNA-Seq to measure translation efficiency (*i.e.*, ribosome footprints normalized by RNA abundance) and found that mRNAs beginning with AUA and AUG did not show a consistent trend in translation efficiency change (Figure 5D). In contrast, AUU-initiated *MT-ND2* mRNA was activated in *NSUN3* KO cells. Thus, f^5^C34 may function in translation initiation in an mRNA-dependent manner.

### τm^5^U34 for UUG, UGG, and AAG codons

Taurine modification (τm^5^U34) is a mt-tRNA-specific modification (Figure 4A and S4B). This modification has been found in 5 mt-tRNAs, mt-tRNA^Leu (UUR)^, mt-tRNA^Trp^, mt-tRNA^Lys^, mt-tRNA^Gln^, and mt-tRNA^Glu^, which decode the NNR codons (where R represents A or G). The MTO1-GTPBP3 complex catalyzes τm^5^U using taurine and 5,10-methylenetetrahydrofolate (CH_2_-THF) ^13,63,72–75^ (Figure 4A and S4B). Among these 5 tRNAs, U34 in mt-tRNA^Lys^, mt-tRNA^Gln^, and mt-tRNA^Glu^ is additionally modified to s^2^U by MTU1 and is ultimately converted to τm^5^s^2^U34 ^73,76–79^ (Figure 4A and S4B). The τm^5^ and s^2^ modifications are mutually independent ^73^.

MitoIP-Thor-Ribo-Seq with *GTPBP3* KO cells showed that the impact of τm^5^U34 modification varies across codons decoded by the five mt-tRNAs (Figures 4B, S4M, and S6A-E). First, τm^5^U34 did not affect ribosome traversal on codons ending in A (UUA, UGA, AAA, CAA, and GAA) (Figures 4B, S4M, and S6A-E), as taurine modification-free tRNA (U or s^2^U) could base-pair with these codons. Second, among the G-ending codons, UUG, UGG, and AAG led to mitoribosome stalling due to GTPBP3 depletion at both the A site and P site (Figures 4B, S4M, S6A-E, and S9A). UUG codons (located at the A site) in *MT-ND2* exemplified the mitoribosome stalling induced by *GTPBP3* KO (Figure S6F). The weak effect on the GAG Glu codon may be associated with the low τm^5^ modification rate on mt-tRNA^Glu^ at the basal level ^63^.

Among the 13 ORFs, *MT-ND6*, in which the mitoribosome footprint distribution was most disrupted (Figure 4D), exhibited mitoribosome arrest at an AAG codon (P-site position) and concomitantly reduced mitoribosome flux downstream (Figure S6G).

During the data analysis, we noted that the mitoribosome occupancy changed beyond the codons recognized by the τm^5^-modified mt-tRNA. We detected greater mitoribosome occupancy at the CAU His codons (Figures 4B, S4M, and S6H). This finding suggested that τm^5^-hypomodified mt-tRNA^Gln^, which typically decodes the CAG and CAA codons, may compete with mt-tRNA^His^ for the CAU codon. Considering that a substantial fraction of mt-tRNA^Gln^ was not fully modified with τm^5^s^2^ at the basal level and only possessed τm^5^ ^60^, GTPBP3 depletion may increase the amount of unmodified U34 in mt-tRNA^Gln^. Since mt-tRNA with U34 could recognize all four bases (U, A, G, and C) at the wobble position (*i.e.*, 4-way wobbling) ^61^, the hypomodified mt-tRNA^Gln^ may have a greater frequency of binding to the nearcognate CAU codon. Thus, ultimately, the CAU codon may require a longer time to be decoded by mt-tRNA^His^ (Figure S9A).

### MELAS-causing mt-tRNA mutation leads to mitoribosome queuing

τm^5^U deficiency has been implicated in diseases such as mitochondrial myopathy, encephalopathy, lactic acidosis, and stroke-like episodes (MELAS). The most frequent mutation associated with MELAS in the mitochondrial genome is A3243G; this site corresponds to the 14th position of mt-tRNA^Leu (UUR) 80–82^ (Figure 5E). Destabilization of the conformation by the mutation ^83,84^ may cause the tRNA to become a poor substrate of the MTO1-GTPBP3 complex and thus reduce its τm^5^U modification rate ^63,85–87^.

We explored the translational dynamics in the myoblasts of a MELAS patient with A3243G ^87,88^ via MitoIP-Thor-Ribo-Seq (Figures 4B, 4H, and S4L-M). Consistent with earlier biochemical work ^89^, we observed the accumulation of mitoribosomes at UUG codons (Figures 4B, S4M, and S6I), as was found in *GTPBP3* KO cells (Figures 4B, S4M, and S6A). Again, the footprint accumulation at both the P and A sites suggested a function for τm^5^U in decoding and translocation. This pausing was represented by a UUG codon in *MT-ND2* (Figure 5F). In stark contrast, no alteration of reads on UUA codons was observed (Figures 4B, S4M, and S6I).

Strikingly, we observed mitoribosome queuing in MELAS patient cells. A metagene plot around UUG codons revealed 2 upstream peaks at 11-12 codon intervals (Figure 5G left), which is the size of a mitoribosome (Figure 1J), indicating the formation of collided mito-disomes and mito-trisomes. We also found a high buildup of mitoribosomes downstream of UUG (Figure 5G left). This may stem from the misincorporation of different amino acids and the generation of difficult peptides held in the exit tunnel of mitoribosomes and/or membrane insertion. Indeed, amino acid misincorporation has been suggested in the A3243G MELAS patient cell line ^88^. Alternatively, this mitoribosome stalling on the out-of-frame stop codon (enriched downstream of UUG codons, Figure S6K) may cause this phenotype due to a frameshift at the UUG codon. An earlier study also detected short truncated proteins in ^35^S-labeling of the newly synthesized proteins ^88^. In sharp contrast, the distribution of mitoribosomes around UUA codons was not affected (Figure 5G right). These data indicated that severe mitoribosome stalling occurs in MELAS patients.

We also noted a bias in footprint distribution toward the 5′ ends of the ORFs in *MT-ND6* and *MT-ND5* (Figure 4H, 5H, and S6J). These data explain the remarkable reduction in the synthesis rate of these proteins reported in previous studies ^88^. *MT-ND6* is the most UUG-rich mRNA, which has been proposed to explain the susceptibility of MT-ND6 protein synthesis to disruption in MELAS ^89^. Our mitoribosome distribution findings suggest another cause for the high sensitivity of *MT-ND6*.

### s^2^U34 for GAG and GAC codons

Among the three mt-tRNAs possessing s^2^U34 (Figure S4B), the GAG codon showed the strongest mitoribosome pauses by MTU1 depletion (Figures 4B, S4M, S6L-N, and S9A), suggesting dysfunction of mt-tRNA^Glu^. In addition, we also detected that the subcognate GAC Asp codon of this tRNA accumulated in mitoribosomes (Figures 4B, S4M, S6O, and S9A), as if s^2^-hypomodified mt-tRNA^Glu^ competed with mt-tRNA^Asp^ for nearcognate GAC codons. *MT-CO1* provided an example of the mitoribosome stalling on GAG and GAC codons (Figure S6P). Given that similar completion and resultant mitoribosome slowdown were not found in τm^5^-deficient cells (Figure S6Q), the maintenance of fidelity was attributed to s^2^ but not τm^5^ on mt-tRNA^Glu^. These data provide another example of decoding fidelity maintenance by mt-tRNA modifications.

### Q34 for UAU codons

Q modification ^60,90^ is found in both cytosolic and mitochondrial tRNAs and is mediated by the QTRT1-QTRT2 complex, a guanine transglycosylase ^91^. In mitochondria, G34-to-Q34 modifications were detected in mt-tRNA^Asn^ (for the AAU and AAC codons), mt-tRNA^His^ (for the CAU and CAC codons), mt-tRNA^Asp^ (for the GAU and GAC codons), and mt-tRNA^Tyr^ (for the UAU and UAC codons) ^60^ (Figures 4A and S4C). MitoIP-Thor-Ribo-Seq of *QTRT1* KO and *QTRT2* KO cells showed weak but significant stalling of mitoribosomes on UAU Tyr codons at both the A and P sites (Figure 4B, S4M, S7, and S9A). This observation was consistent with our earlier reports of conventional Ribo-Seq in *QTRT2* KO cells ^60^.

### t^6^A37 for AAA and AAG codons

In addition to modification at the 34th position, we studied a modification at the 37th position, 3′ adjacent to the anticodon (Figure 4A). Generally, the modification of this position is thought to contribute to accurate codon recognition and the prevention of frameshift errors ^92,93^. We investigated t^6^A modification at this position ^60^. This modification, catalyzed by OSGEPL1 ^64,94,95^, is found in mt-tRNAs responsible for the following ANN codons: mt-tRNA^Lys^, mt-tRNA^Asn^, mt-tRNA^Thr^, mt-tRNA^Ile^, and mt-tRNA^Ser (AGY)^ (where Y represents U or C) (Figures 4A and S4D).

Among the codons decoded by t^6^A-modified mt-tRNAs, OSGEPL1 depletion led to mitoribosome stalling on the AAG and AAA codons (Figures 4B, S4M, and S8A-F). These effects on Lys codons may be due to reduced aminoacylation of mt-tRNA^Lys^ ^64,95^. An AAG codon in *MT-CO1* provided a remarkable example of the mitoribosome pause (Figure S8G). In addition, the distribution of the mitoribosome footprint was biased toward the 5′ end in *MT-ND6* (Figure 4G); AAG at the 21st codon (at the P-site position) induced mitoribosome stalling and a subsequent reduction in mitoribosome traversal downstream (Figure S8H). Strikingly, the same codon position was similarly sensitive to the loss of τm^5^U34 modification (Figure S6G).

Moreover, CCU Pro codons exhibited increased mitoribosome occupancy upon OSGEPL1 deletion (Figure 4B, S4M, and S8I). Given that t^6^A37 strengthens the base pairing between U36 (in the tRNA position) and A1 (as the first position of the codon) ^96,97^, the loss of t^6^A could lead to the misrecognition of subcognate codons with U1, C1, or G1. The accumulation of mitoribosomes on CCU codons may be caused by the competition of hypomodified mt-tRNA^Thr^ for ACN codons with mt-tRNA^Pro^ for CCU codons (Figure S9B). Amino acid misincorporation reported upon OSGEPL1 depletion ^95^ may stem from such competition between mt-tRNAs.

Overall, MitoIP-Thor-Ribo-Seq allowed us to evaluate the significance of all the modifications at the 34th position and a well-conserved modification at the 37th position in mt-RNA and to reveal the unexpected specificity of those modifications for codon and fidelity maintenance functions (Figure S9).

### Investigation of mitoribosome collision

Along with the mt-tRNA modifications that facilitate smooth elongation, the programmed slowdown of mitoribosome traversal may benefit protein synthesis. Given the importance of cotranslational insertion of the transmembrane domain ^98–104^ and cotranslational complex assembly ^105^, mitoribosome stalling may provide time for nascent peptides to target the membrane and interact with assembly factors.

To survey ribosome stall sites at high resolution, longer ribosome footprints generated by colliding cytoribosomes have been analyzed (disome profiling) ^106–116^. Since we omitted the sucrose density gradient to isolate the 55S mito-monosome and instead implemented MitoIP, our method could capture the longer footprint that potentially originated from colliding mitoribosomes (Figure 6A). Here, we focused on 50- to 80-nt long RNA fragments generated by RNase treatment of isolated mitochondrial fractions (Figure 6A) and found footprints of the expected size (Figure S10A) (MitoIP-Thor-Disome-Seq).

**Figure 6.**
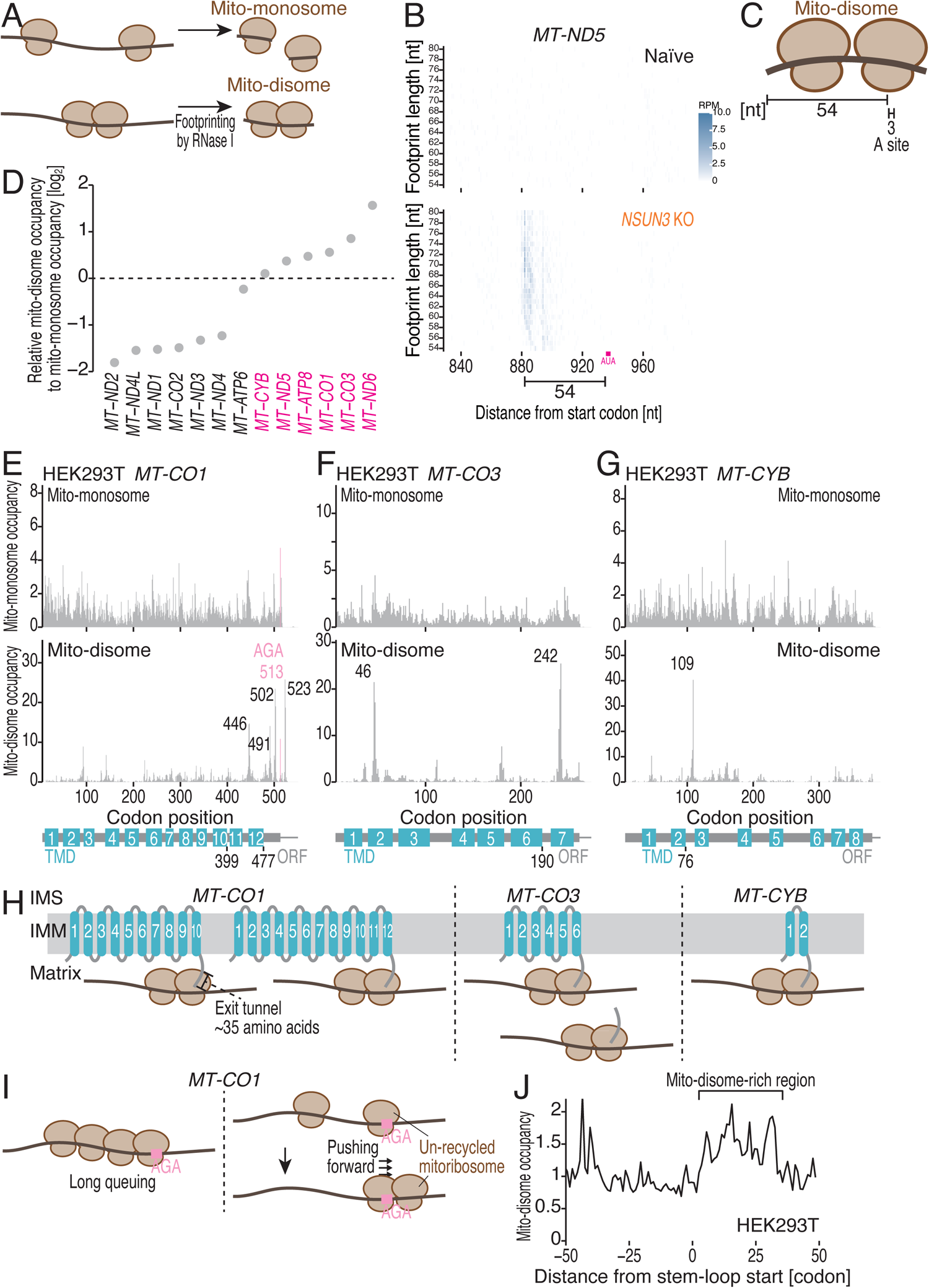
Mitoribosome pause sites revealed by MitoIP-Thor-Disome-Seq. (A) Schematic of mitochondrial disome profiling. (B) Plots of the 5′ ends of the mito-disome footprints around AUA-AUA (at 312-313; the first nucleotide of the start codon was set to 0) dicodons in *MT-NT5* in the indicated cells. The color scale indicates the read abundance. (C) Schematic representation of the relative position of the A site in the leading paused mitoribosome for mito-disome footprints. (D) Relative abundance of mito-disome footprints to mito-monosome footprints on the indicated transcripts. The data include the reads from overlapping ORF regions in *MT-ATP8*/*MT-ATP6* and *MT-ND4L*/*MT-ND4*. (E-G) Distribution of mito-monosome and mito-disome footprints from naïve HEK293 cells along the indicated transcripts. The A-site position (in the case of mito-disomes, the leading paused mitoribosome) of each read is indicated. The AGA stop codon is highlighted. (H) Schematic of the configurations of the mito-disome pause sites found in *MT-CO1*, *MT-CO3*, and *MT-CYB* with cotranslational membrane insertion of their nascent peptides. (I) Schematic of mitoribosome collision around the noncanonical AGA stop codon associated with inefficient termination and recycling reactions. (J) Metagene plots for mito-disome footprints around the RNA regions with secondary structures. See also Figure S10 and Table S1.

To precisely assess the position of leading, stalled mitoribosomes in the mito-disome footprints, we harnessed mitoribosomes stalled on AUA-AUA dicodons (codon position 312-313, with the start codon at position 0) on *MT-ND5* by *NSUN3* KO (Figure 5B). In naïve HEK293T cells, mito-disome footprints were quite sparse in this region (Figure 6B top). However, NSUN3 depletion dramatically increased the number of reads around the AUA-AUA dicodon (Figure 6B bottom). Considering the distance between the 5′ ends of the read and the AUA-AUA dicodon, the A-site offset was determined to be 54 nt for the mito-disome footprints (Figure 6C). This A-site assignment of reads enabled us to assess the position of stalled mitoribosomes at codon resolution. In addition to *MT-ND5* (Figure S10B), *MT-ND1* exhibited a striking peak of mito-disome footprints on the AUA-AUA dicodon (Figure S10C). Since the mito-disome fraction did not include normally elongating mitoribosomes, MitoIP-Thor-Disome-Seq allowed further sensitive detection of mitoribosome pause sites in *NSUN3* KO (Figure S10D-E).

MitoIP-Thor-Disome-Seq identified mitoribosome collision events that occurred naturally, even in the absence of mt-tRNA modification defects. We detected differences in the occurrence of mito-disome and mito-monosome footprints across mRNAs (Figure 6D). Among the mito-disome-rich transcripts, we observed a sharp accumulation of mito-disome footprints at specific positions (Figures 6E-G and S10F-G). Considering that the mitoribosome exit tunnel accommodates amino acids 30-40 of the nascent peptide ^104^, the mito-disome pause sites were assigned to the position where the completed transmembrane domain (TMD) was exposed to the solvent. For example, position 446 on *MT-CO1* allowed the synthesis of TMD10, with the C-terminal end facing the matrix (Figure 6E and 6H). A similar topology between TMD synthesis and mito-disome pause sites was found for *MT-CO1* TMD12, *MT-CO3* TMD6, *MT-CYB* TMD2, *MT-ND6* TMD1, and *MT-ATP8* TMD1 (Figures 6E-H and S10F-H). Notably, due to the overlapping ORFs of *MT-ATP8* and *MT-ATP6* in the bicistronic transcript, we could not clearly distinguish whether the leading mitoribosome in the mito-disome (codon 65 in the *MT-ATP8* position) (Figure S10G) was on *MT-ATP8* or *MT-ATP6*, although the trailing mitoribosome should be translated with *MT-ATP8* (Figure S10H).

On *MT-CO1,* in which mitoribosome collision frequently occurs, we detected significant mito-disomes on the noncanonical AGA stop codon ^41^ (position 513), suggesting inefficient termination/recycling of this noncanonical stop codon (Figure 6E). Furthermore, mitoribosomes may form a long queue, as indicated by monitoring mito-disomes ∼10 codons upstream (position 502) and ∼20 codons upstream (position 491) (Figure 6E). Thus, the noncanonical AGA stop codon serves as the source of mitoribosome queuing (Figure 6I left).

In addition to the ORF, we found mito-disome formation on the 3′ UTR of *MT-CO1* (position 523) (Figures 6E and S10I). The leading paused mitoribosome was located 10 codons downstream of the stop codon, the size of a single mitoribosome, whereas the trailing mitoribosome was on the ORF. The disome positions were not associated with in-frame or out-of-frame stop codons in the 3′ UTR (Figure S10I), excluding frameshifts or readthroughs and subsequent pauses at the next stop codon. As the unrecycled cytoribosome could be “pushed” by translating ribosomes and form disomes until the trailing ribosomes reach the stop codons ^110,111^, this mito-disome may be produced by the same process (Figure 6I, right).

Importantly, the mitoribosome collisions found in HEK293T cells were also observed in HAP1 cells and HeLa cells at the same codons (Figure S10J-N), suggesting the commonality of the regulation across cell types. The mitoribosome collision may be associated with mRNA secondary structures, which were probed by mitoDMS-MaP-Seq ^117^ (Figure 6J). Overall, MitoIP-Thor-Disome-Seq offers a means to monitor elongation slowing, termination, and recycling events in mitochondrial protein synthesis.

## Discussion

The improved mitochondrial ribosome profiling methods developed here illustrated a complex landscape of *in organello* translation. By comparing cytosolic translation kinetics assessed by our recent work ^26^, which used Ribo-Calibration and cytoribosome run-off assays for standard Ribo-Seq, we found that mitochondrial translation kinetics were strikingly different from those of cytosolic translation measures in the same HEK293 cells: the elongation rate was ∼10 times slower [0.52 codon/s in mitochondria (this study) vs. 4.1 codon/s in the cytoplasm ^26^]; the initiation rate was even slower than the elongation rate in both translation systems but more extreme in mitochondria [440 s/event in mitochondria (this study) vs. ∼22 s/event in the cytoplasm ^26^]. These inefficiencies may be compensated for by the high expression of mitochondrial mRNAs (Figure S10O). A recent study by Churchman’s group ^55,118^ reached a similar conclusion regarding a slow translation rate in mitochondria. Ultimately, the protein supply from the mitochondrial matrix and the cytosol should be balanced for the maintenance of proper proteostasis ^16,119,120^.

Moreover, our study revealed that mitochondrial translation kinetics could vary among cell types (Figure 2). Our study illustrated the coordination of the translation initiation and elongation rates; both kinetics were faster in C2C12 cells than in HEK293 cells (Figure 2). These cell type-dependent kinetics may explain the high vulnerability of some tissues to mitochondrial disease ^121,122^. It is not surprising that kinetic regulation may be induced by the extra- and intracellular cues, as indicated by our recent work ^123^.

The function of τm^5^U34 in mitochondrial translation suggested by this study was largely consistent with an earlier report in which conventional MitoRibo-Seq was applied to *MTO1* KO cells and A3243G MELAS patient fibroblasts ^13^. Our improved methods further extended our understanding of the role of τm^5^U34’s role in mitochondrial translation fidelity (*i.e.*, suppressing the competition of mt-tRNA^Gln^ with mt-tRNA^Asp^ for the CAU Asp codon). Moreover, our data also revealed long queues of mitoribosomes upstream of UUG codons in MELAS patient cells. These data highlighted the utility of MitoIP-Thor-Ribo-Seq for detecting pathogenesis-related translational events.

Our data revealed the functional similarities and differences in specific modifications between mitochondrial and cytosolic tRNAs. Q34 is also found in cytosolic tRNAs (ct-tRNA) decoding the same set of codons (*i.e.*, ct-tRNA^Asn^, ct-tRNA^His^, ct-tRNA^Asp^, and ct-tRNA^Tyr^) ^124,125^. Standard Ribo-Seq of cells cultured in Q-depleted media showed slow decoding of all the U-terminal codons (such as AAU, CAU, GAU, and UAU) ^90^, but this decrease was more pronounced than that we found for mitochondria (Figures 4B, S4M, and S7). This suggested that the two translational systems rely on the Q modification to different extents.

s^2^U34 was found in the same tRNA species in the cytosol and mitochondria (*i.e.*, ct-tRNA^Lys^, ct-tRNA^Gln^, and ct-tRNA^Glu^). These base positions are simultaneously modified with 5-metoxycarbonylmethyl (mcm^5^) and ultimately form mcm^5^s^2^U34 ^61^. Thus, mcm^5^ in cytosolic tRNA could be a counterpart of τm^5^ in the mitochondria. In contrast, the loss of mcm^5^ and s^2^ in cytosolic tRNA does not impact the wobble pairing codons (G-ending) of AAG, CAG, or GAG ^126–128^, which are critically impacted by mitochondrial τm^5^ and s^2^ (Figures 4B, S4M, and S6). Rather, those modifications facilitate the decoding of Watson–Crick pairing (A-ending) codons of AAA, CAA, and GAA ^126–128^. Nonetheless, mcm^5^ in ct-tRNA^Gln^ and τm^5^ in mt-tRNA^Gln^ can suppress the miscoding of subcognate CAU codons (Figure S6H) ^127^. Our study illuminated the distinction in the functionality of similar (mcm^5^ vs. τm^5^) and identical (s^2^) modifications at the U34 position in the two translation systems.

The role of t^6^A37 modification in cytosolic tRNAs, including the same five ct-tRNAs as in their mitochondrial counterparts, ct-tRNA^Met^, and ct-tRNA^Arg (AGR)^ (where R represents A or G), has been investigated via yeast Ribo-Seq ^128,129^. The data indicated that t^6^A37 has pleiotropic effects: it may increase the elongation rate of U-ending codons (ANU), whereas it has the opposite effect on C-ending codons (ANC). Our data showed more limited effects on translation; AAG and AAA codon decoding were only facilitated by t^6^A37 (Figures 4B, S4M, and S8). Another important activity of t^6^A37 in cytosolic tRNAs is fidelity maintenance by suppressing leaky scanning, stop codon readthrough, and frameshifting ^130^. However, the molecular basis of this function is attributed to the prevention of the competition of t^6^A37-modified elongator tRNA^Met^ with the nonmodified initiator tRNA^Met^ ^130^. On the other hand, in mitochondria, the same mt-tRNA^Met^ has both translation initiation and elongation functions and is not modified by t^6^A37 ^60^. Thus, mitochondrial t^6^A37 should not have such a role. Indeed, our data did not show evidence of stop codon readthrough (Figure S8J).

Through the development of MitoIP-Thor-Disome-Seq, we identified mitoribosome arrest sites at single-codon resolution. Earlier work biochemically predicted mitoribosome pauses in *MT-CO1* mRNA at the ∼212 and ∼280 codon positions, which engage a complex IV assembly intermediate termed MITRAC ^105^. However, the corresponding sites were not identified in our mito-disome profiling or even mito-monosome profiling (Figures 6E and S10I). The lack of corresponding pause sites was also reported in an earlier MitoRibo-Seq study ^12^. At present, we cannot explain this discrepancy. Future studies, such as the isolation of MITRAC and associated mitoribosomes and subsequent mito-monosome/disome profiling, will uncover the pause sites accompanied by the complex assembly.

## Limitations

Since lifetime translation rounds are a relatively new parameter, a validation method is lacking. Future work is needed to support our measurements.

## Acknowledgments

We thank all the members of the Iwasaki laboratory for constructive discussion and technical assistance. We also thank Dr. Nono Takeuchi-Tomita, Dr. Layla Kawarada, and Dr. Shunpei Okada for their fruitful comments and technical advice on this work. MELAS myoblasts were kind gifts from Dr. Brendan J. Battersby. S.I. was supported by the Ministry of Education, Culture, Sports, Science and Technology (MEXT) (JP20H05784 and JP24H02307), the Japan Society for the Promotion of Science (JSPS) (JP23H02415 and JP23H00095), the Japan Agency for Medical Research and Development (AMED) (JP20gm1410001), the Gushinkai Foundation, and RIKEN (Pioneering project “Biology of Intracellular Environments”). Y.S. was supported by MEXT (JP21H05734 and JP23H04268), JSPS (JP21K15023 and JP23K05648), AMED (JP23gm6910005), and RIKEN (Pioneering project “Biology of Intracellular Environments” and Incentive Research Projects). T.W. was supported by The Graduate School of Frontier Sciences, The University of Tokyo (C2205 and C2306)] and JSPS (JP23KJ0444). Tsutomu S. was supported by the Japan Science and Technology Agency (JST) (JPMJER2002). A portion of the DNA libraries was sequenced at the National Institute of Genetics and at the Vincent J. Coates Genomics Sequencing Laboratory at QB3 Genomics, UC Berkeley, Berkeley, CA (RRID: SCR_022170), which is supported by the NIH S10 OD018174 Instrumentation Grant. The computations were supported by the supercomputer HOKUSAI SailingShip in RIKEN. We are grateful to the Support Unit for Bio-Material Analysis, RIKEN CBS Research Resources Division, for Sanger sequencing and cell sorting. T.W. was a recipient of fellowships from JSPS (DC2), JST SPRING (JPMJSP2108), and the ANRI. H.T. was a JSPS postdoctoral fellow (PD).

## Author contributions

Conceptualization, T.W., M.M., H.Y., and S.I.;

Methodology, T.W., M.M., H.Y., Kotaro T., H.T., Y.S., and S.I.;

Formal analysis, T.W., H.Y., Kotaro T., H.T., Y.S., and S.I.;

Investigation, T.W., M.M., H.Y., Kotaro T., H.T., and Y.S.;

Resources, H.T., Kazuhito T., T.C., A.N., Takeo S., F.Y.W., and Tsutomu S.;

Writing – Original Draft, S.I.;

Writing – Review & Editing, T.W., M.M., H.Y., Kotaro T., H.T., Kazuhito T., T.C., A.N., Takeo S., F.Y.W., Y.S., Tsutomu S., and S.I.;

Visualization, T.W. and S.I.;

Supervision, F.Y.W., Kazuhito T., T.C., A.N., Takeo S., Y.S., Tsutomu S., and S.I.; Project administration, S.I.;

Funding Acquisition, T.W., Y.S., and S.I.

## Conflicts of interest

S.I. is a member of the *Scientific Reports* editorial board.

## Materials and Methods

### Cell lines

*NSUN3* KO (lines #1 and #2), *GTPBP3* KO, *QTRT1* KO, *QTRT2* KO, and *OSGEPL1* KO (line #2) HEK293T cells were generated previously ^60,62–64,68^. Wild-type or A3243G-mutated myoblasts were previously derived from a muscle sample from a MELAS patient and clonally selected ^87,88^. Along with the cell lines above, naïve HEK293T cells, HEK293 cells [American Type Culture Collection (ATCC), CRL-1573], naïve C2C12 cells (ATCC, CRL-1772), and naïve HeLa cells (RIKEN BioResource Research Center) were cultured at 37°C with 5% CO_2_ in the media described in Table S1. An e-Myco VALiD Mycoplasma PCR Detection Kit (iNtRON Biotechnology) was used to confirm that the culture was free of *Mycoplasma*.

#### Animals

Male wild-type mice and heart-specific *Nsun3* KO mice were acquired previously ^71^. Mice were housed at 25°C with a 12-h:12-h light-dark cycle. The experiment was performed at 8 weeks of age. After sacrifice, the cardiac muscles were dissected, immediately flash-frozen in liquid nitrogen, and stored at −80°C. All animal experiments were authorized and approved by the animal ethics committee of Kumamoto University (approval ID: A2021-012R3 and A2023-011).

#### Generation and validation of MTU1 KO cells

*MTU1* KO HEK293 cells were generated using the CRISPR-Cas9 system. A guide RNA sequence was selected to target exon 1 of *MTU1* and cloned into LentiCRISPR v2 (a gift from Feng Zhang, Addgene plasmid # 52961; http://n2t.net/addgene:52961; RRID: Addgene_52961) ^131^. The targeting sequence was 5′-GCTGTCCACGCCGCCGGGACA-3′. Cells were infected with lentivirus carrying the above gRNA and Cas9 gene and selected with puromycin for 3 d. Single colonies were selected using a cloning disc. Total cell lysates from candidate cell lines were subjected to Western blotting. Total RNA from each candidate cell line was digested to single nucleosides using nuclease P1 (Fujifilm Wako Pure Chemical Corporation) and bacterial alkaline phosphatase (TaKaRa), followed by mass spectrometry analysis, as previously described ^77^. The MTU1-mediated mitochondrial tRNA τm^5^s^2^U (*m/z* 398.0686) modification was detected using a Q Exactive mass spectrometer (Thermo Fisher Scientific) in full mass scan and positive ion mode. ms^2^i^6^A (*m/z* 382.1543), a mitochondrial tRNA-specific modification, was used as a control for mitochondrial RNA loading.

#### Generation and validation of mtIF3 KO cells

*mtIF3* KO cells were generated via CRISPR-mediated genome editing according to a previously described protocol ^132^. Two guide RNA sequences were selected to target exon 1 of *mtIF3* as described previously (5′-GCAAUAGGGGACAACUGUGC-3′ and 5′-GCAGAGUAUCAGCUCAUGAC-3′) ^59^ and cloned into the pL-CRISPR.EFS.GFP (a gift from Benjamin Ebert, Addgene plasmid # 57818; http://n2t.net/addgene:57818; RRID: Addgene_57818) and pL-CRISPR.EFS.tRFP (a gift from Benjamin Ebert, Addgene plasmid # 57819; http://n2t.net/addgene:57819; RRID: Addgene_57819) ^133^, respectively. After verification by Sanger sequencing, both plasmids containing sgRNAs were transfected into naïve HEK293 cells by Lipofectamine 3000 (Thermo Fisher Scientific) as instructed by the manufacturer. At 48 h posttransfection, RFP and BFP dual-positive populations of cells were sorted by flow cytometry. After cultivation until visible colonies were observed, single clones were expanded in a 96-well dish. *mtIF3* KO clones were confirmed by Sanger sequencing of genomic DNA (Figure S3K) and Western blotting (Figure S3L).

#### MitoIP-Ribo-Seq, MitoIP-Thor-Ribo-Seq, and MitoIP-Thor-Disome-Seq

##### Lysate preparation

Cells were cultured in 15-cm dishes, washed with ice-cold PBS, and lysed with 1200 µl of hypotonic buffer (10 mM HEPES-KOH pH 7.5, 10 mM KCl, 1.5 mM MgCl_2_, 1 mM DTT, 100 µg/ml cycloheximide, and 100 µg/ml chloramphenicol). The lysate was incubated with anti-TOM22 antibody-conjugated beads from a Mitochondria Isolation Kit, Human (Miltenyi Biotec) or a Mitochondria Isolation Kit, mouse tissue (Miltenyi Biotec) for 60 min at 4°C. The subsequent wash and elution steps were performed with wash buffer (10 mM HEPES-KOH pH 7.5, 10 mM KCl, 15 mM MgCl_2_, 1 mM DTT, 100 µg/ml cycloheximide, and 100 µg/ml chloramphenicol) according to the manufacturer’s instructions. The eluted mitochondrial fraction was pelleted by centrifugation at 7,000 × g for 10 min at 4°C and resuspended in modified lysis buffer (20 mM Tris-HCl pH 7.5, 150 mM NaCl, 15 mM MgCl_2_, 1 mM DTT, 1% Triton X-100, 100 µg/ml cycloheximide, 100 µg/ml chloramphenicol, and 2.5 U/ml TURBO DNase). After clarification by centrifugation at 20,000 × g for 10 min at 4°C, the supernatant was subjected to library preparation.

For the mitoribosome run-off assay, cells were pretreated with 100 µg/ml retapamulin for 30, 60, 90, 120 s, or 30 min and then incubated with 100 µg/ml chloramphenicol and 100 µg/ml cycloheximide to arrest translation elongation. Subsequently, cell lysis and mitochondrial purification were performed as described above, and 100 µg/ml retapamulin was added to all the solutions.

##### Mouse cardiac muscle

The frozen myocardium was pulverized with 600 µl of liquid nitrogen-frozen hypotonic buffer using a Multi-beads Shocker (Yasui Kikai) at 3000 rpm for 30 sec for 2 cycles. The lysate was clarified by an EASYstrainer (Greiner Bio-One), and mitochondria were isolated as described above.

##### Library preparation; MitoIP-Ribo-Seq

The lysate was incubated with 2 U of RNase I (LGC Biosearch Technologies, low), 10 U of RNase I (high), 0.12 U of RNase A/T1 (Thermo Fisher Scientific, low), or 0.6 U of RNase A/T1 (high) in a 50 µl reaction (scaled up by the modified lysis buffer) at 25°C for 45 min. Then, for Ribo-FilterOut ^26^, the mitoribosomes were pelleted by sucrose cushion and resuspended in EDTA lysis buffer [20 mM Tris-HCl pH 7.5, 150 mM NaCl, 15 mM MgCl_2_, 1 mM DTT, 5 mM EDTA, 1% Triton X-100, and 20 U/ml SUPERase•In RNase inhibitor (Thermo Fisher Scientific)]. The solution was loaded onto an Amicon Ultra 0.5 ml Ultracel 100K centrifugal filter (Millipore) and centrifuged at 14,000 × *g* at 4°C for 10 min ^26^. The RNA from the flowthrough was purified with TRIzol LS (Thermo Fisher Scientific) and a Direct-zol RNA MicroPrep Kit (Zymo Research) and separated on a 15% UREA PAGE gel. Fragments ranging from 17 to 50 nucleotides (nt) were excised from the gel. Subsequent library generation followed an earlier report ^28^: RNA fragments were dephosphorylated and ligated with linkers; rRNA depletion was conducted with a Ribo-Zero Gold rRNA Removal Kit (Human/Mouse/Rat) (Illumina, accompanied by a TruSeq Stranded Total RNA Kit); cDNA was reverse-transcribed, circularized by ligation, and PCR-amplified. Sequencing of the DNA libraries was performed on a HiSeq 4000 platform in 50-bp single-read mode.

To test translation inhibitors, chloramphenicol in hypotonic buffer, wash buffer, and modified lysis buffer was replaced with capreomycin, blasticidin S, or tigecycline.

A size-exclusion gel-filtration spin column was used as described previously ^28^.

##### Library preparation; MitoIP-Thor-Ribo-Seq and MitoIP-Thor-Disome-Seq

Library preparation was conducted as described above with modifications. The lysate was incubated with 40 U of RNase I (LGC Biosearch Technologies) in a 50 µl reaction (brought up to volume with the modified lysis buffer) at 25°C for 45 min. Fragments ranging from 17 to 50 nt for monosomes and from 50 to 80 nt for disomes were collected. Then, the libraries were prepared as previously reported ^27^. From linker-ligated RNA fragments, rRNAs were depleted with Human Ribo-Seq riboPOOL (siTOOLs Biotech) or Mouse/Rat Ribo-Seq riboPOOL (siTOOLs Biotech). Then, an oligonucleotide was hybridized to the T7 promoter region of the linker to generate the dsDNA T7 promoter. Complementary RNAs were transcribed with a T7-Scribe Standard RNA IVT kit (CELLSCRIPT). After ligation of the second linker, cDNA was reverse-transcribed and PCR-amplified. DNA libraries were sequenced on a HiSeq X Ten platform or a NovaSeq X Plus platform in 150-bp paired-end mode or a NovaSeq 6000 platform in 100-bp single-end mode. The experimental design can be found in Table S1.

#### Standard Ribo-Seq, RNA-Seq, and Ribo-Calibration

##### Lysate preparation

Approximately 1 × 10^7^ cells in a 10-cm dish were lysed with 600 µl of lysis buffer (20 mM Tris-HCl pH 7.5, 150 mM NaCl, 5 mM MgCl_2_, 1% Triton X-100, 1 mM DTT, 100 µg/ml cycloheximide, and 100 µg/ml chloramphenicol) and treated with 25 U/ml TURBO DNase (Thermo Fisher Scientific) ^28^. Then, the lysate was clarified by centrifugation at 20,000 × g for 10 min at 4°C.

##### Library preparation; standard Ribo-Seq and RNA-Seq

Standard Ribo-Seq library preparation was performed according to a previously described protocol ^26,28^. After RNase digestion and sucrose cushion ultracentrifugation, Ribo-FilterOut was conducted. Then, ribosome footprints ranging from 17-34 nt were gel-excised.

For RNA-Seq, total RNA was extracted from the same lysate used for Ribo-Seq with TRIzol LS (Thermo Fisher Scientific) and a Direct-zol RNA MicroPrep Kit (Zymo Research). Four hundred nanograms of total RNA were used for a SEQuoia Express Standard RNA Library Prep Kit (Bio-Rad).

For both library preparations, rRNA depletion was performed with a Ribo-Zero Gold rRNA Removal Kit (Human/Mouse/Rat) (Illumina, accompanied by a TruSeq Stranded Total RNA Kit). Libraries were sequenced on a HiSeq X Ten platform in 150-bp paired-end mode.

##### Library preparation; Ribo-Calibration

Ribo-Calibration library preparation was conducted as described previously ^26^. The lysate of C2C12 cells containing 10 µg of total RNA was adjusted to a volume of 340 µl using the lysis buffer. Then, a 10 µl spike-in mixture of Rluc-disome and Fluc-trisome was added to the lysate to make a 350 µl lysate solution.

For Ribo-Seq, 290 µl of the lysate solution was incubated with 10 µl of 10 U/µl RNase I (Lucigen) for 45 min at 25°C. After sucrose cushion ultracentrifugation and Ribo-FilterOut ^26^, fragments ranging from 17 to 34 nt were collected. Then, the libraries were prepared as described above (see the “*Library preparation; MitoIP-Thor-Ribo-Seq and MitoIP-Thor-Disome-Seq*” section). rRNA depletion was conducted with Mouse/Rat Ribo-Seq riboPOOL (siTOOLs Biotech). Libraries were sequenced on a NovaSeq 6000 platform in 100-bp single-end mode.

For RNA-Seq, the remaining 50 µl of the lysate solution was directly subjected to RNA purification with TRIzol LS reagent (Thermo Fisher Scientific) and a Direct-zol RNA MicroPrep Kit (Zymo Research). Five hundred nanograms of total RNA were used for a SEQuoia Express Standard RNA Library Prep Kit (Bio-Rad). rRNA was depleted with Mouse/Rat riboPOOL (siTOOLs Biotech). Libraries were sequenced on a HiSeq X Ten platform in 150-bp paired-end mode.

#### Data analysis

Deep sequencing data processing was performed as described previously ^134,135^ with modifications. For paired-end sequencing, fastp (version 0.21.0) ^136^ was used for base correction and read 1 was processed for downstream analysis. Read quality filtering and adapter sequence removal were conducted by fastp. All reads were aligned to noncoding RNAs (rRNAs, tRNAs, mt-rRNAs, mt-tRNAs, snRNAs, snoRNAs, and miRNAs), and the remaining reads were then mapped to human or mouse nuclear genomes (human, hg38; mouse, mm10) and a custom database of mitochondrial transcript sequences using STAR (version 2.7.0a) ^137^. For MitoIP-Thor-Ribo-Seq and MitoIP-Thor-Disome-Seq, duplicated reads were suppressed according to the unique molecular index (UMI) in the linkers using UMItools (version 1.1.2) ^138^.

Ribosomal A-site offsets for each footprint length were empirically estimated (Table S1). For the analysis of the first 5 codons, the dedicated A-site offsets were estimated separately considering the short or absent 5′ UTRs in the mitochondrial mRNAs (except for *MT-ATP6* and *MT-ND4*) (Table S1).

The relative entropy of *H_j_* for read length *j* was defined as follows:

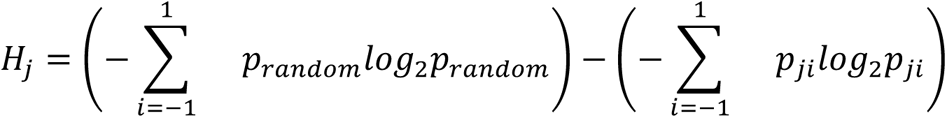

where *p_ji_* is the fraction of mitoribosomes with footprint length *j* at frame *i* and *p_random_* is 0.333. Then, the periodicity score (*P*) was defined as follows:

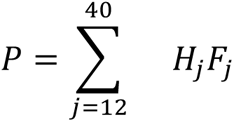

where *F_j_* represents the fraction of mitoribosomes with footprint length *j*.

To calculate the mitoribosome occupancy, the reads of each codon were normalized to the average reads per codon of the mitochondrial transcript. We omitted overlapping ORF regions in the bicistronic transcripts from the calculation (except where indicated in the figure panel legends).

The polarity score was defined as previously reported ^65^.

To calculate translation efficiency, a generalized linear model in the DESeq2 package ^139^ was used based on the relative enrichments of standard Ribo-Seq reads in ORFs over those in RNA-Seq. Reads assigned to the first and last 5 amino acids were excluded from this analysis.

For the mitoribosome run-off assay, smoothed metagene profiles were plotted from the mitochondrial transcripts, excluding *MT-ATP6* and *MT-ND4*. The mean read density from 100 to 200 codons for HEK293 cells and from 150 to 250 codons for C2C12 cells was set to 1. The number of mitoribosomes and the half-life of the mitochondrial transcripts were determined from Ribo-Calibration and BRIC-Seq data as described previously ^26^.

The translation initiation rate (*I*) for mitochondrial transcript *k* was defined as follows:

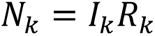

where *N_k_* and *R_k_* represent the number of mitoribosomes and the residence time of the loaded mitoribosome on transcript *k,* respectively. Since the loaded mitoribosome undergoes an elongation reaction on the ORF length *L_k_* of transcript *k*, *R_k_* was defined as follows:

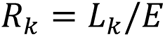

where *E* represents the global mitochondrial elongation rate, which was determined to be 0.515 codon/s for HEK293 cells and 1.013 codon/s for C2C12 cells by the mitoribosome run-off assay. Thus,

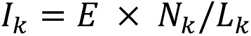

The lifetime translation round (*R*) for mitochondrial transcript *k* was defined as follows:

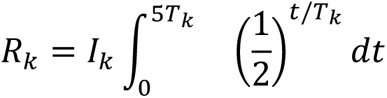

where *T_k_* represents the half-life of transcript *k*. The ribosome loading frequency was calculated as 5*T_k_* when ∼97% of the mRNA is predicted to be degraded.

To define mitochondrial translation initiation sites from retapamulin-assisted MitoIP-Thor-Ribo-Seq data, the A-site assigned reads at each nucleotide were normalized by the average reads per nucleotide of the transcript. The ROC curve was generated using the score at 13 annotated translation initiation sites as true positives. Then, the threshold was set at 99% specificity (1% false positive rate), which identified the peaks. We omitted the P sites within 3 nt upstream and 28 nt downstream of the annotated translation initiation sites. If the sites overlapped within 3 nt, the site with the highest value was chosen. We searched for possible AUG, AUA, and AUU codons and their subcognate codons within a 5-nt range (the peak position and 4 nt upstream). If those codons were not found, the 3 nt sequence upstream of the peak position was considered.

### Sucrose density gradient

The cell lysate (see the “*Standard Ribo-Seq and RNA-Seq*” section) or purified mitochondrial fraction (see the “*MitoIP-Ribo-Seq, MitoIP-Thor-Ribo-Seq, and MitoIP-Thor-Disome-Seq*” section) was loaded onto a 10-50% sucrose gradient in SDG buffer (20 mM Tris-HCl pH 7.5, 150 mM NaCl, 5 mM MgCl_2_, 1 mM DTT, and 100 µg/ml cycloheximide) and ultracentrifuged at 38,000 rpm and 4°C for 4 h by a Himac CP80WX ultracentrifuge (Hitachi) with a P40ST rotor (Hitachi). The gradients were fractionated with continuous measurement of absorbance at 260 nm using a TRIAX flow cell (BioComp) and micro collector (ATTO).

#### On-gel mito-FUNCAT

On-gel mito-FUNCAT was performed as described previously ^50,140^. HEK293 cells were cultured in a 6-well plate in methionine-free medium [DMEM, high glucose, no glutamine, no methionine, no cystine (Thermo Fisher Scientific), supplemented with 48 µg/ml L-cysteine and 4.08 mM L-alanyl-L-glutamine] supplemented with 50 µM homopropargylglycine (HPG) (Jena Bioscience) and 100 mg/ml anisomycin (Alomone Labs, A-520) for 3 h before cell harvest. During HPG labeling, the cells were treated with the following compounds: tetracycline (5 μg/ml), chloramphenicol (100 μg/ml), fusidic acid (300 μg/ml), spectinomycin (300 μg/ml), hygromycin (300 μg/ml), G418 (150 μg/ml), kanamycin (60 μg/ml), streptomycin (300 μg/ml), capreomycin (300 μg/ml), blasticidin S (30 μg/ml), tigecycline (300 μg/ml), linezolid (300 μg/ml), thiostrepton (300 μg/ml), or retapamulin (100 μg/ml). An equivalent volume of solvent was used as a control for each compound. After being washed with ice-cold PBS, the cells were lysed with mito-FUNCAT lysis buffer (20 mM Tris-HCl pH 7.5, 150 mM NaCl, 5 mM MgCl_2_, and 1% Triton X-100) and clarified by centrifugation at 20,000 × g for 10 min at 4°C. The HPG-labeled proteins in the supernatants were labeled with 50 μM IRdye800CW Azide (LI-COR Biosciences) with a Click-it Cell Reaction Buffer Kit (Thermo Fisher Scientific) according to the manufacturer’s instructions. The reactants were run on SDS– PAGE gels, and the specific signals were detected by an Odyssey CLx (LI-COR Biosciences) in the IR 800-nm channel. For standardization, the gels were stained with GelCode Blue Safe Protein Stain (Thermo Fisher Scientific) to monitor the protein input using the IR 700-nm channel. The images were quantified with Image Studio (LI-COR Biosciences, version 5.2).

#### Western blotting

For Figures S1B and S3L, anti-MRPS22 (Thermo Fisher Scientific, PA5-52249, 1:1000), anti-MRPL45 (Thermo Fisher Scientific, PA5-54784, 1:1000), anti-RPL10A (Abcam, ab174318, 1:1000), anti-RPS17 (Abcam, ab1128671, 1:1000), anti-GAPDH (Cell Signaling Technology [CST], 2118, 1:1000), anti-TOMM20 (CST, 42406, 1:1000), anti-mtIF3 (Proteintech, 14219-1-AP, 1:1000), and anti-β-actin (MEDICAL & BIOLOGICAL LABORATORIES [MBL], M177-3, 1:1000) were used as primary antibodies, and IRDye800CW anti-rabbit IgG (LI-COR Biosciences, 926-32211, 1:10000) was used as the secondary antibody. Images were acquired with an Odyssey CLx (LI-COR Biosciences) and Image Studio (LI-COR Biosciences, version 5.2).

For Figure S4E, anti-MTU1 (Sigma–Aldrich, AV48819, 1:2000) and anti-VDAC (CST, 4866, 1:2000) antibodies were used as primary antibodies, and anti-rabbit IgG and HRP-linked antibody (CST, 7074, 1:5000) was used as a secondary antibody. Antibodies were diluted with 5% skim milk (nacalai tesque, 31149-75) in PBS (Thermo Fisher Scientific, 10010023). Images were acquired with Fusion Solo S (VILBER).

#### Material availability

The materials generated in this study will be distributed upon request. There are restrictions to availability due to a material transfer agreement (MTA).

#### Data availability

The results of the MitoIP-Ribo-Seq, MitoIP-Thor-Ribo-Seq, standard Ribo-Seq, and RNA-Seq (GEO: GSE237154) obtained in this study have been deposited in the National Center for Biotechnology Information (NCBI) database. This study also used the reported data for standard Ribo-Seq, RNA-Seq, and BRIC-Seq (GSE233555 and GSE233374) ^26^.

#### Code availability

The key custom scripts used in this study are available at Zenodo (https://zenodo.org, DOI: 10.1101/2023.07.19.549812).

---

**Figure S1.**
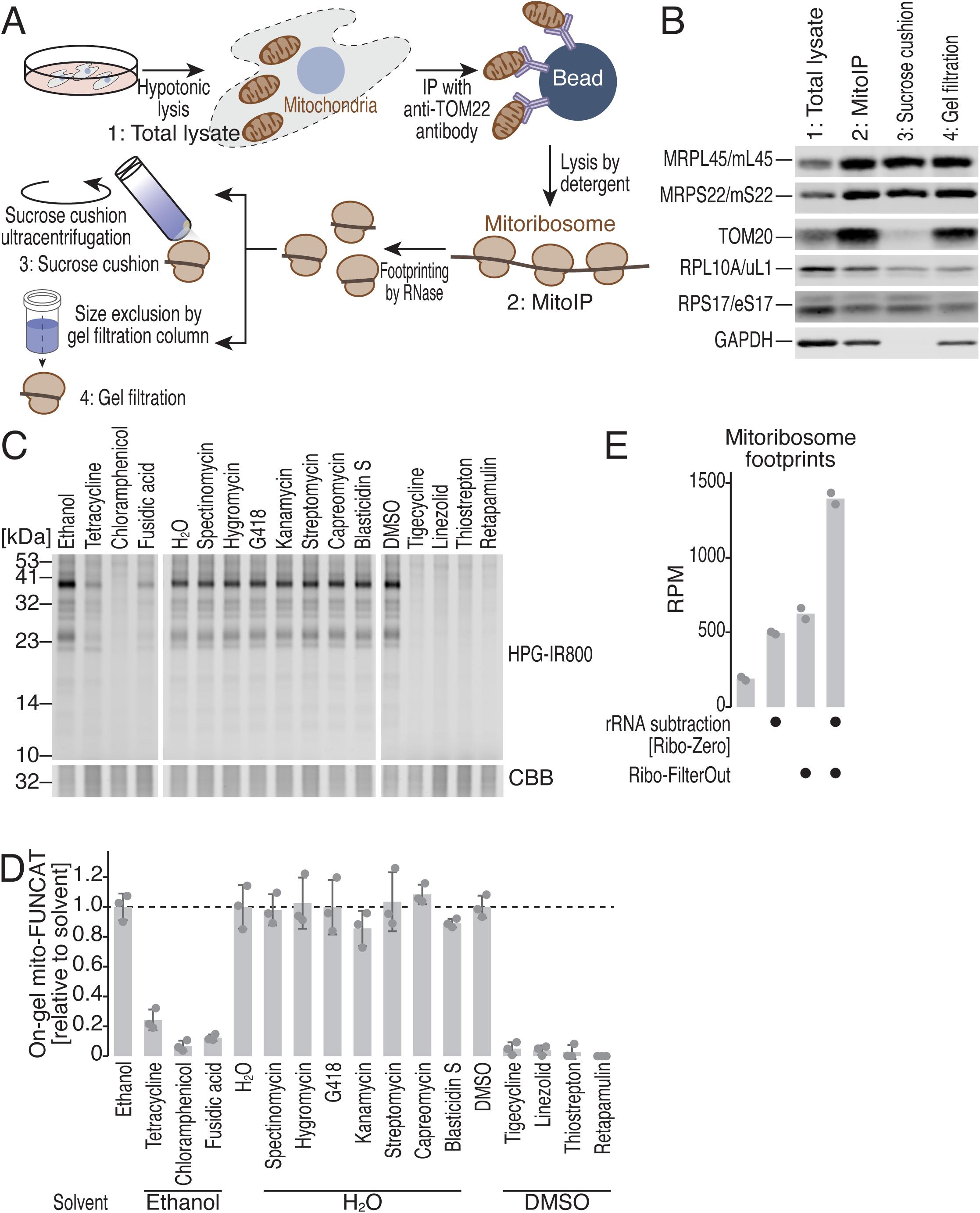
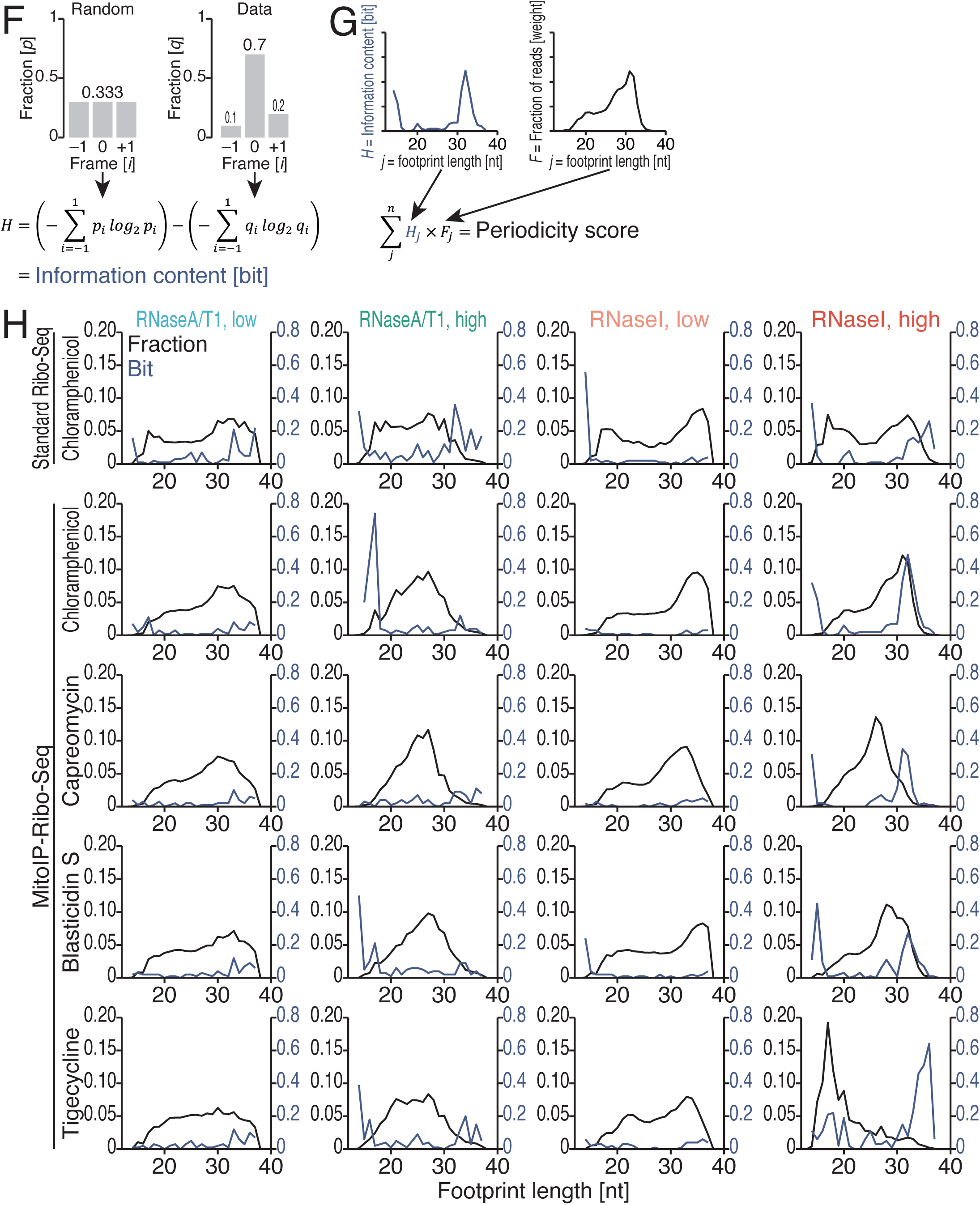
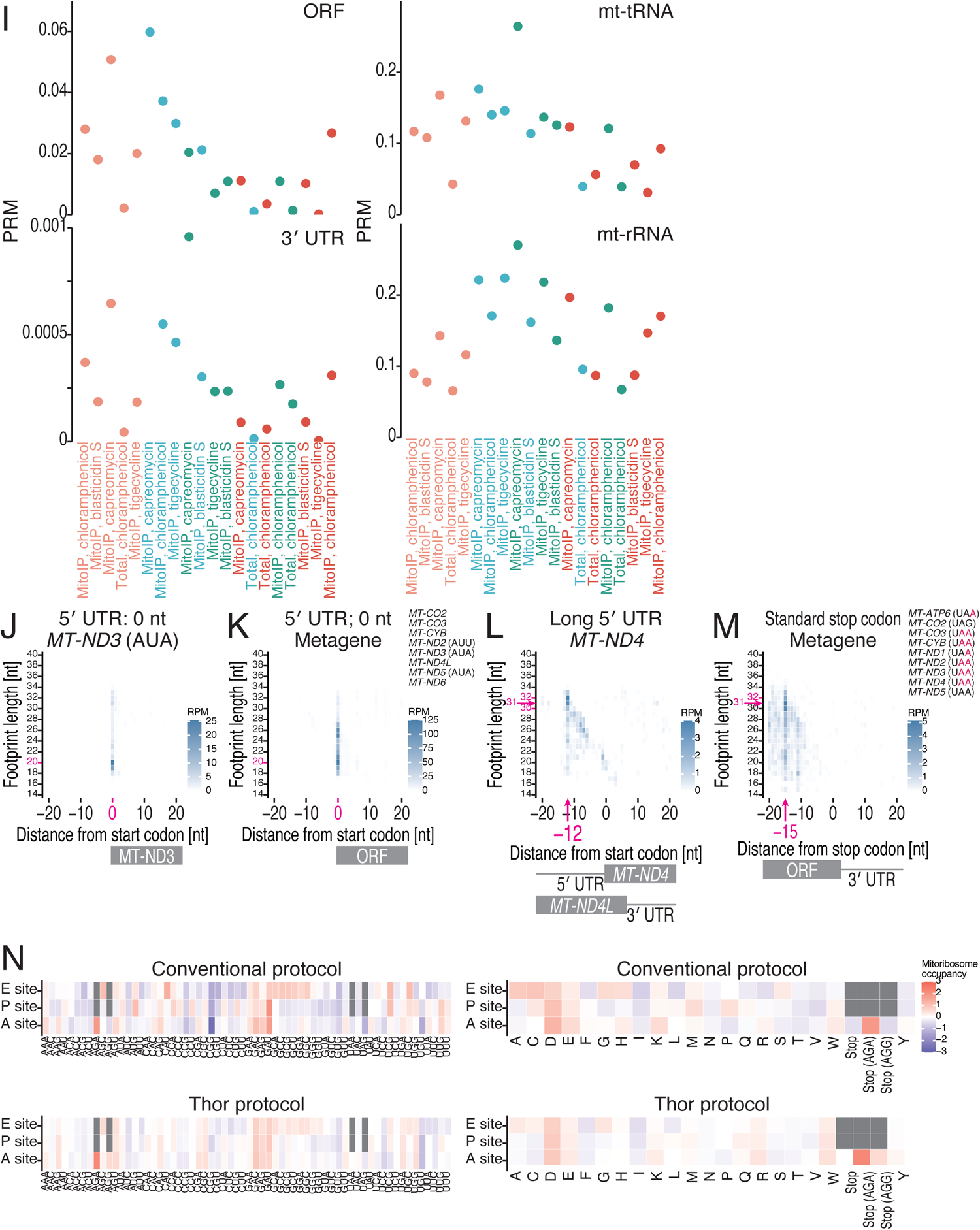
Optimization of MitoIP-Ribo-Seq, related to Figure 1. (A) Schematic of sampling for Western blotting in B. (B) Western blotting of MRPL45/mL45, MRPS22/mS22, TOM20, RPL10A/uL1, RPS17/eS17, and GAPDH to evaluate mitoribosome isolation. The RNase I-treated (0.2 U) lysate (or input) was subjected to sucrose cushion ultracentrifugation or a size-exclusion gel-filtration spin column. (C and D) On-gel mito-FUNCAT was used to test the effect of the indicated bacterial ribosome inhibitors on mitochondrial translation (C). CBB staining of the total protein was used for the loading control. The means (bars), s.d.s (errors), and individual replicates (n = 3, points) are shown in D. (E) The combination of the conventional rRNA subtraction method (Ribo-Zero) with Ribo-FilterOut in standard Ribo-Seq. Reads mapped to mitochondrial transcripts were analyzed. The means (bars) and individual replicates (n = 2, points) are shown. (F and G) Schematic of the calculation of information content (F) and the definition of the periodicity score (G). (H) The fraction of mitoribosome footprints and information content along the read length for the indicated conditions. (I) The read abundance of the indicated RNA regions in the indicated conditions of MitoIP-Ribo-Seq. RPM, reads per million mapped reads. (J-L) Plots of the 5′ ends of the mitoribosome footprint along the length around the start codon (the first nucleotide of the start codon was set to 0) for the indicated transcripts. The color scale indicates the read abundance. (M) Metagene plots for the 5′ ends of mitoribosome footprints around stop codons (the first nucleotide of the stop codon was set to 0). *MT-ATP8* and *MT-ND4L* were excluded from the analysis because their ORF stop codons overlapped with those of *MT-ATP6* and *MT-ND4,* respectively. *MT-CO1* and *MT-ND6* were also excluded from the analysis since they possess noncanonical stop codons (AGA and AGG). (N) Mitoribosome occupancy at the A site, P site, or E site for MitoIP-Ribo-Seq (top) and MitoIP-Thor-Ribo-Seq (bottom). Data for the codon sequence (left) and amino acid sequence (right) are shown. The color scale indicates the mitoribosome occupancy.

**Figure S2.**
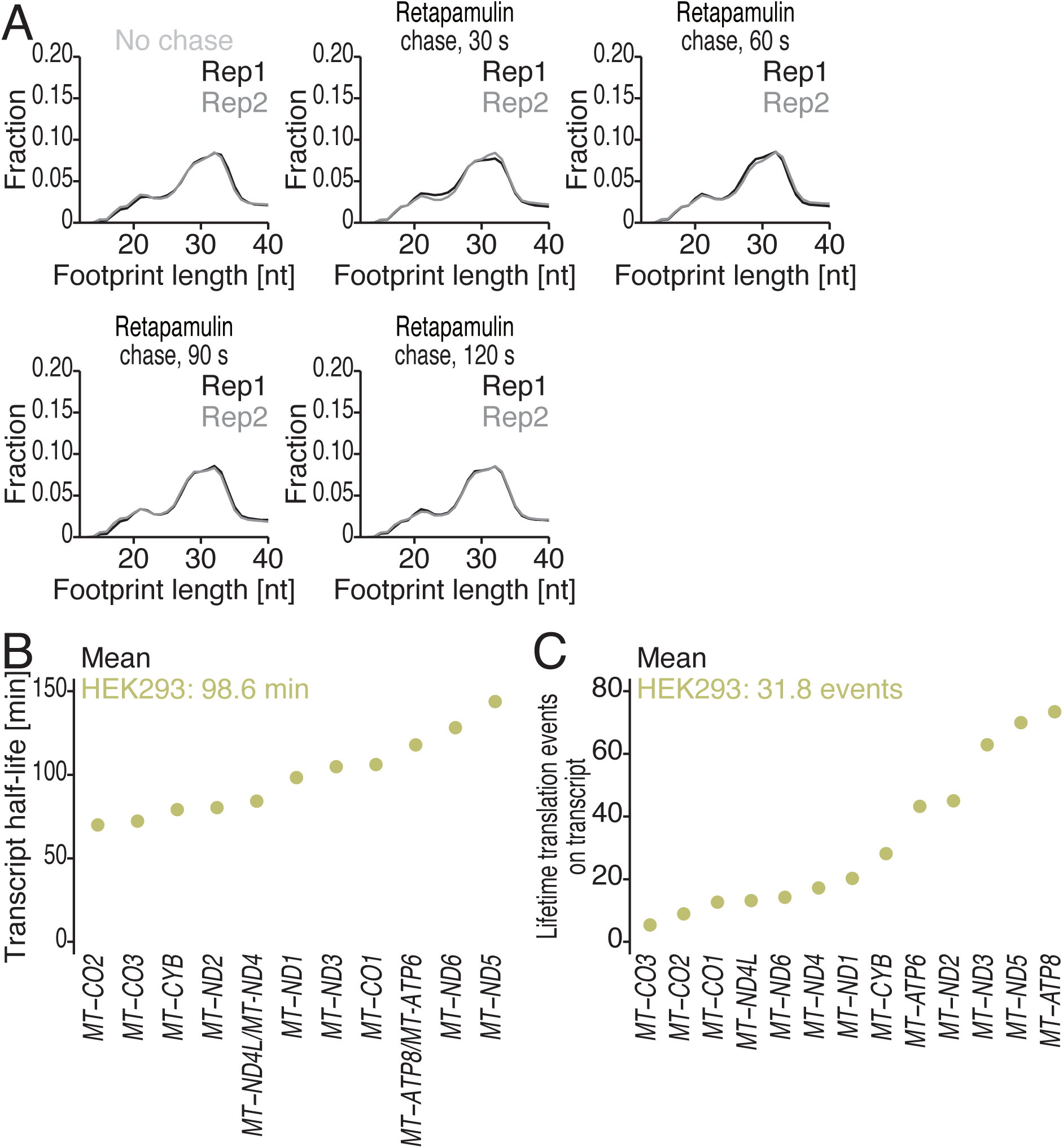
Characterization of mitoribosome run-off assay, related to Figure 2. (A) The fraction of mitoribosome footprints along the read length for the indicated conditions. (B) The transcript half-life of the indicated transcripts, determined by BRIC-Seq ^26^. (C) The lifetime translation rounds before transcript decay of the indicated transcripts.

**Figure S3.**
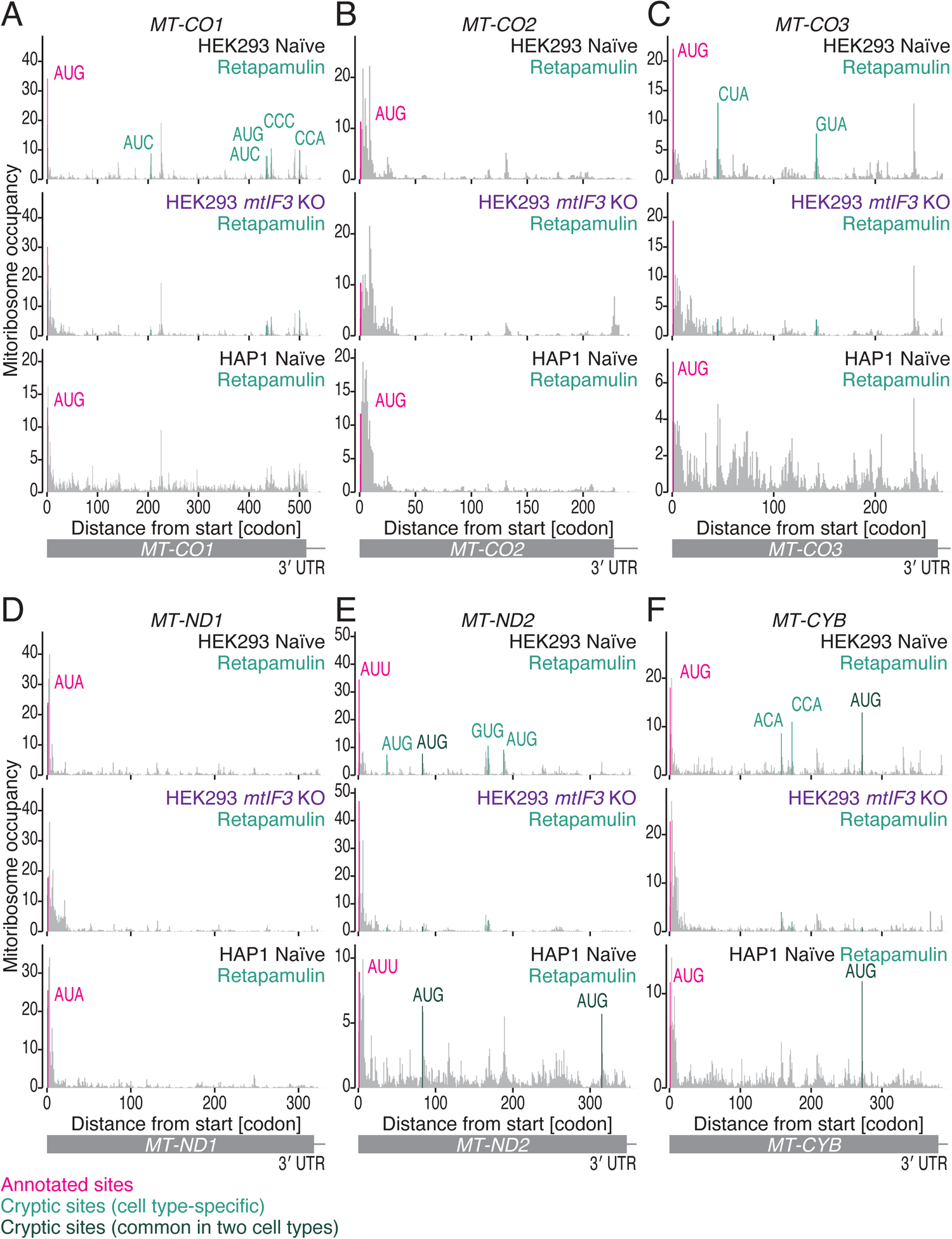
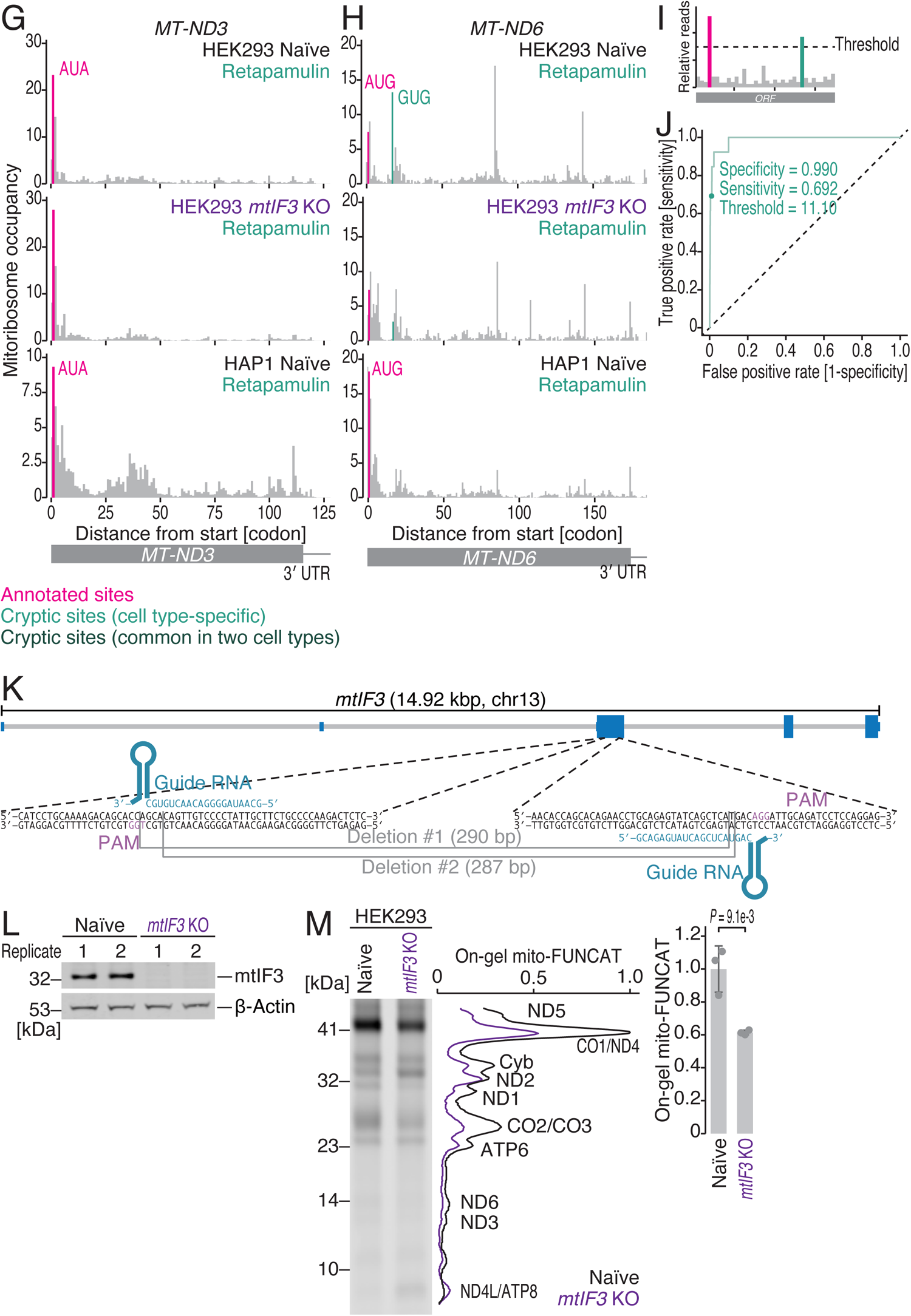
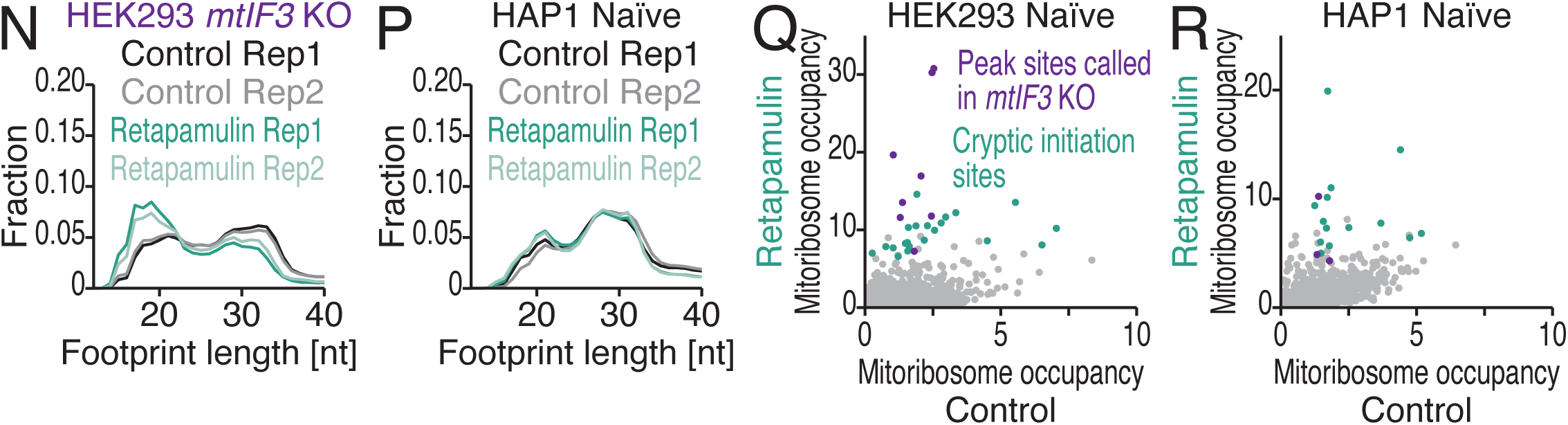
Characterization of retapamulin-assisted MitoIP-Thor-Ribo-Seq data, related to Figure 3. (A-H) Distribution of mitoribosome footprints from retapamulin-treated and control cells along the indicated transcripts. The A-site position of each read is indicated. Codon sequences corresponding to the P site are highlighted. Magenta, annotated initiation sites; peacock green, cryptic initiation sites found in a cell type-specific manner; and dark green, cryptic initiation sites found in both cell lines. See Table S3 for the full list. (I) Schematic of peak calling with the threshold setting. (J) ROC curve to determine the threshold for peak calling. The relative read counts at the annotated translation initiation sites in HEK293 cells were considered true positives. The cutoff point used for this analysis is highlighted. (K) Schematics of guide RNAs (gRNAs) designed for CRISPR-Cas9-mediated gene knock for *mtIF3* and deleted region for the isolated cell line. (L) The loss of mtIF3 protein in *mtIF3* KO cells was validated by Western blotting. Each lane represents a biological replicate. β-Actin was detected as a loading control. (M) On-gel mito-FUNCAT was used to test the effect of mtIF3 deletion. The signal intensities along the lanes are shown in the middle. The means (bars), s.d.s (errors), and individual replicates (n = 3, points) are shown on the right. (N and P) The fraction of mitoribosome footprints along the read length for the indicated conditions. (Q and R) Comparison of mitoribosome occupancy in retapamulin treatment and that in control condition for the indicated cell lines. Cryptic translation initiation sites and peak sites in *mtIF3* KO cells are highlighted.

**Figure S4.**
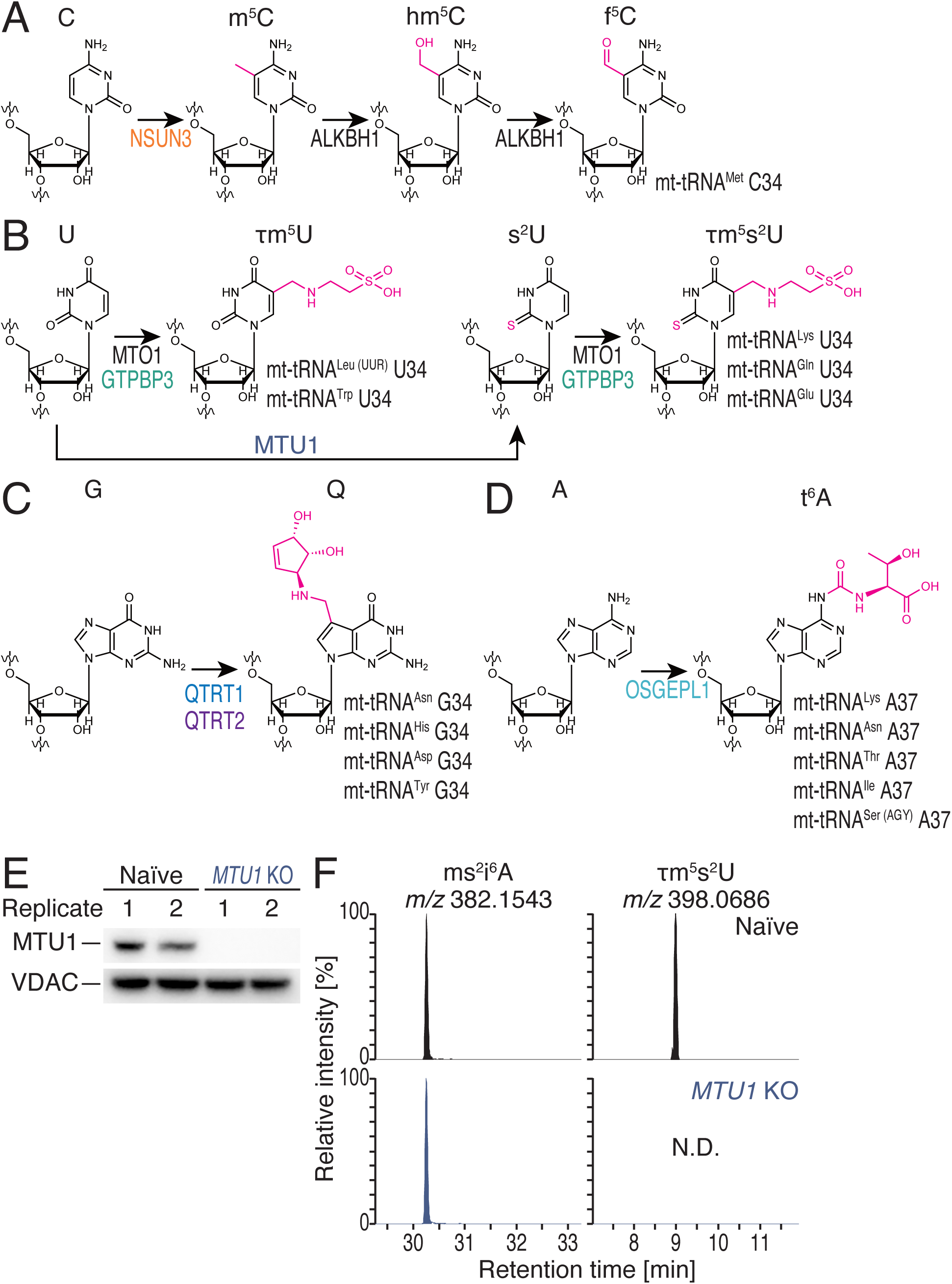
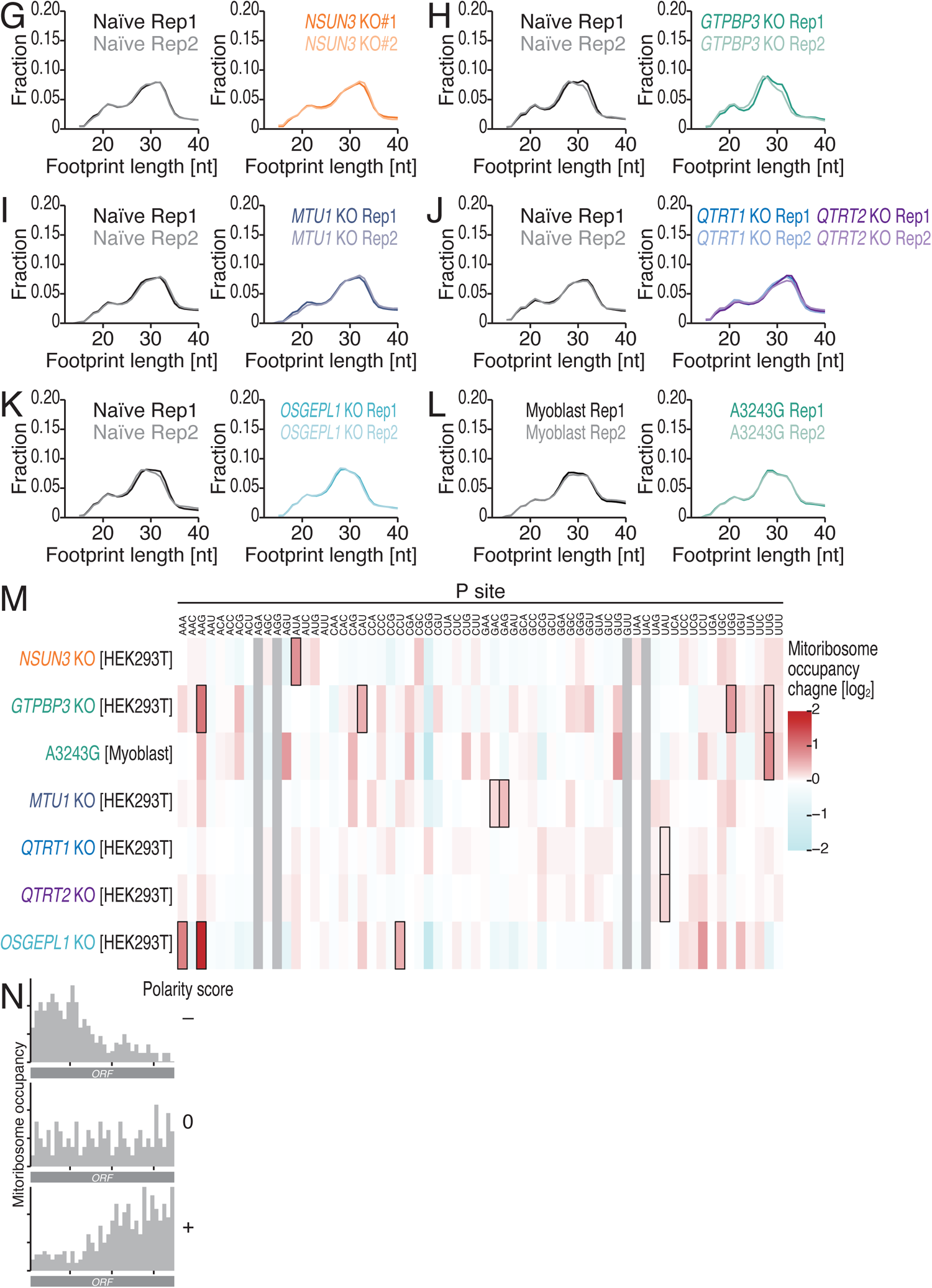
Characterization of MitoIP-Thor-Ribo-Seq data for mt-tRNA modification-deficient cell lines, related to Figure 4. (A) Schematic of f^5^C biosynthesis on mt-tRNA^Met^. (B) Schematic of τm^5^U34 biosynthesis on mt-tRNA^Leu^ ^(UUR)^ and mt-tRNA^Trp^ and τm^5^s^2^U34 biosynthesis on mt-tRNA^Lys^, mt-tRNA^Gln^, and mt-tRNA^Glu^. (C) Schematic of Q34 biosynthesis on mt-tRNA^Asn^, mt-tRNA^His^, mt-tRNA^Asp^, and mt-tRNA^Tyr^. (D) Schematic of t^6^A37 biosynthesis on mt-tRNA^Lys^, mt-tRNA^Asn^, mt-tRNA^Thr^, mt-tRNA^Ile^, and mt-tRNA^Ser^ ^(AGY)^. (E) The loss of MTU1 protein in *MTU1* KO cells was validated by Western blotting. Each lane represents a biological replicate. VDAC, a mitochondrial membrane protein, was detected as a loading control. (F) The loss of mitochondrial tRNA 2-thiolation was investigated with mass spectrometry. τm^5^s^2^U modification disappeared in the total RNA purified from *MTU1* KO cells, whereas ms^2^i^6^A, a mitochondrial tRNA-specific modification, remained intact. (G-L) The fraction of mitoribosome footprints along the read length for the indicated conditions. (M) A heatmap for the mitoribosome occupancy changes at P-site codons in the indicated cell lines. The color scale indicates mitoribosome occupancy change. The remarkable mitoribosome occupancy increases found in the mutant cells are highlighted in black rectangles. (N) Schematic of the polarity score of mitoribosome footprints.

**Figure S5.**
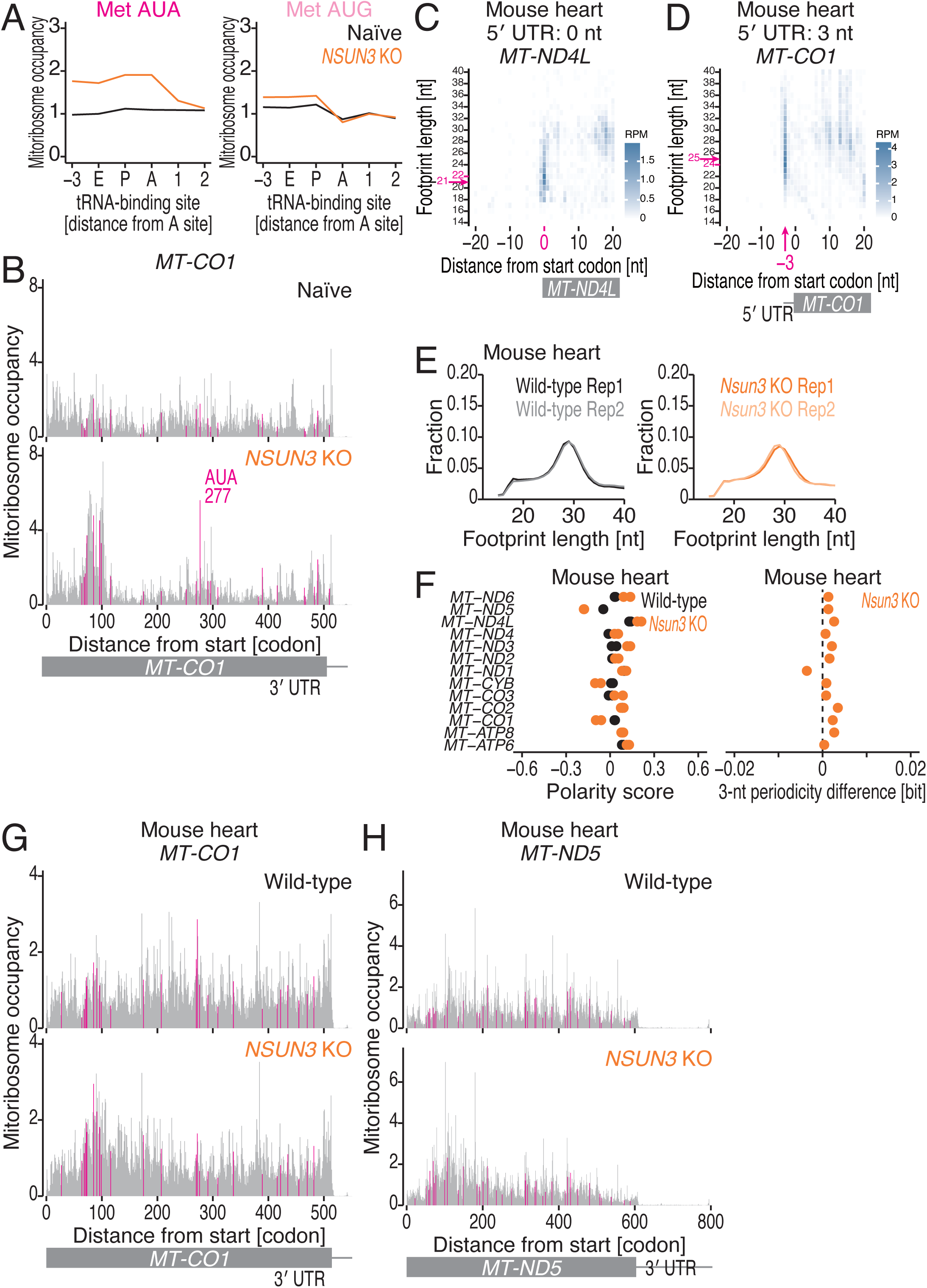
Characterization of MitoIP-Thor-Ribo-Seq data for the heart of *Nsun3* KO mouse, related to Figure 5. (A) Mitoribosome occupancy of naïve and *NSUN3* KO cells for the indicated codons. The scores at each codon position (relative to the A site) are shown. (B) Distribution of mitoribosome footprints from naïve and *NSUN3* KO cells along the indicated transcripts. The A-site position of each read is indicated. AUA codons are highlighted. (C and D) Plots for the 5′ ends of mitoribosome footprints from mouse heart around start codons (the first nucleotide of the start codon was set to 0) for the indicated transcripts. The color scale indicates the read abundance. (E) The fraction of mitoribosome footprints along the read length for the indicated conditions. (F) Polarity scores of the indicated ORFs in the indicated samples (left). The information content for the 3-nt periodicity of the reads from the indicated ORFs was calculated and then the changes by the gene knockout were calculated (right). For the polarity scores, the data points from two replicates are shown individually. For the 3-nt periodicity difference, the means of two replicates are shown. (G and H) Distribution of mitoribosome footprints from the hearts of wild-type and *Nsun*3 KO mice along the indicated transcripts. The A-site position of each read is indicated. AUA codons are highlighted.

**Figure S6.**
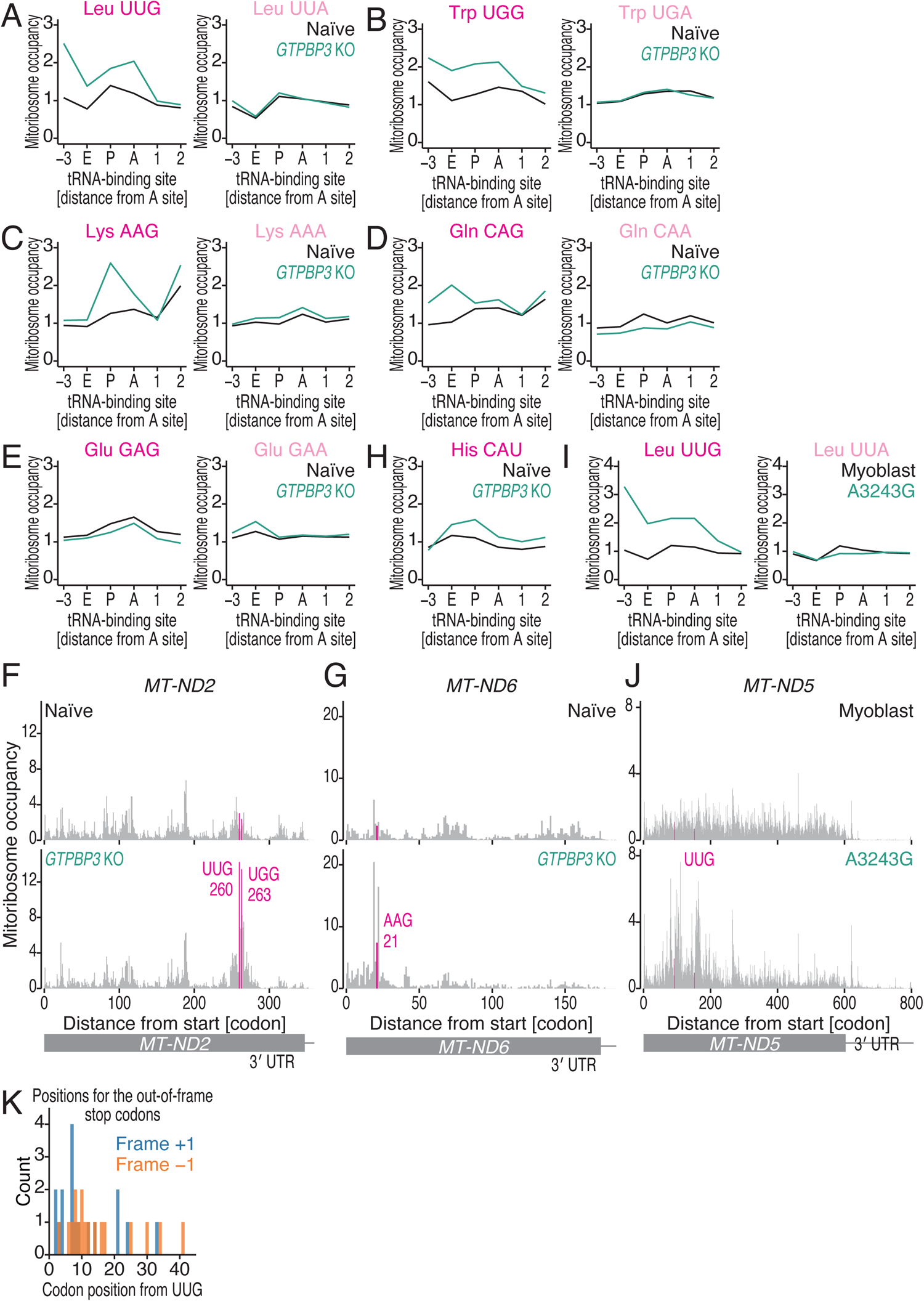
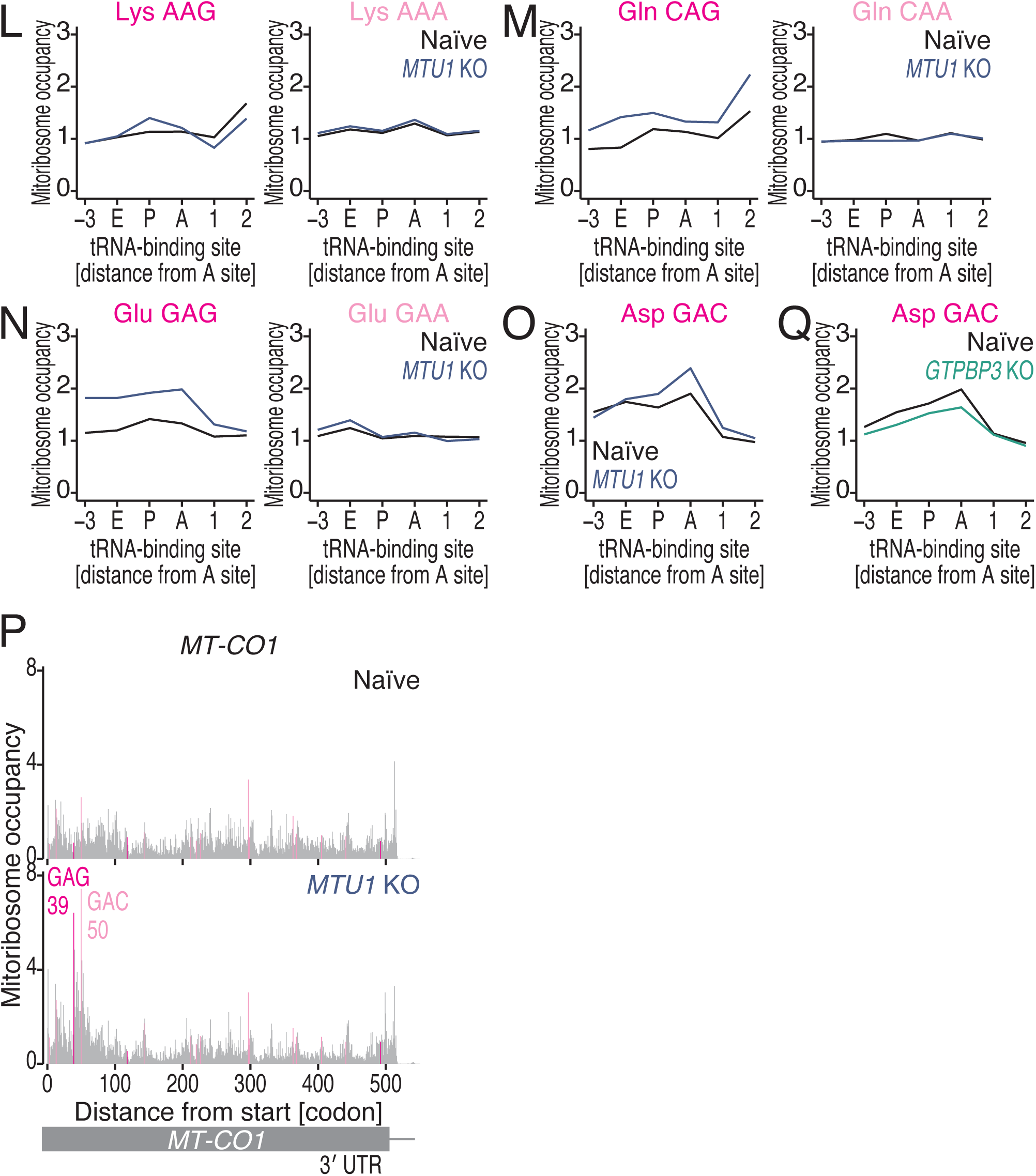
Characterization of MitoIP-Thor-Ribo-Seq data for *GTPBP3* KO cells, A3243G myoblasts, and *MTU1* KO cells, related to Figure 4. (A-E and H) Mitoribosome occupancy of naïve and *GTPBP3* KO cells for the indicated codons. The scores at each codon position (relative to the A site) are shown. (F and G) Distribution of mitoribosome footprints from naïve and *GTPBP3* KO cells along the indicated transcripts. The A-site position of each read is indicated. UUG, UGG, and AAG codons are highlighted. (I) Mitoribosome occupancy of naïve and A3243G myoblasts for the indicated codons. The scores at each codon position (relative to the A site) are shown. (J) Distribution of mitoribosome footprints from naïve and A3243G myoblasts along the indicated transcripts. The A-site position of each read is indicated. UUG codons are highlighted. (K) The position of the out-of-frame stop codons downstream of the UUG codons. (L-O and Q) Mitoribosome occupancy of the indicated codons in naïve and *MTU1* KO cells. The scores at each codon position (relative to the A site) are shown. (P) Distribution of mitoribosome footprints from naïve and *MTU1* KO cells along the indicated transcripts. The A-site position of each read is indicated. GAG and GAC codons are highlighted.

**Figure S7.**
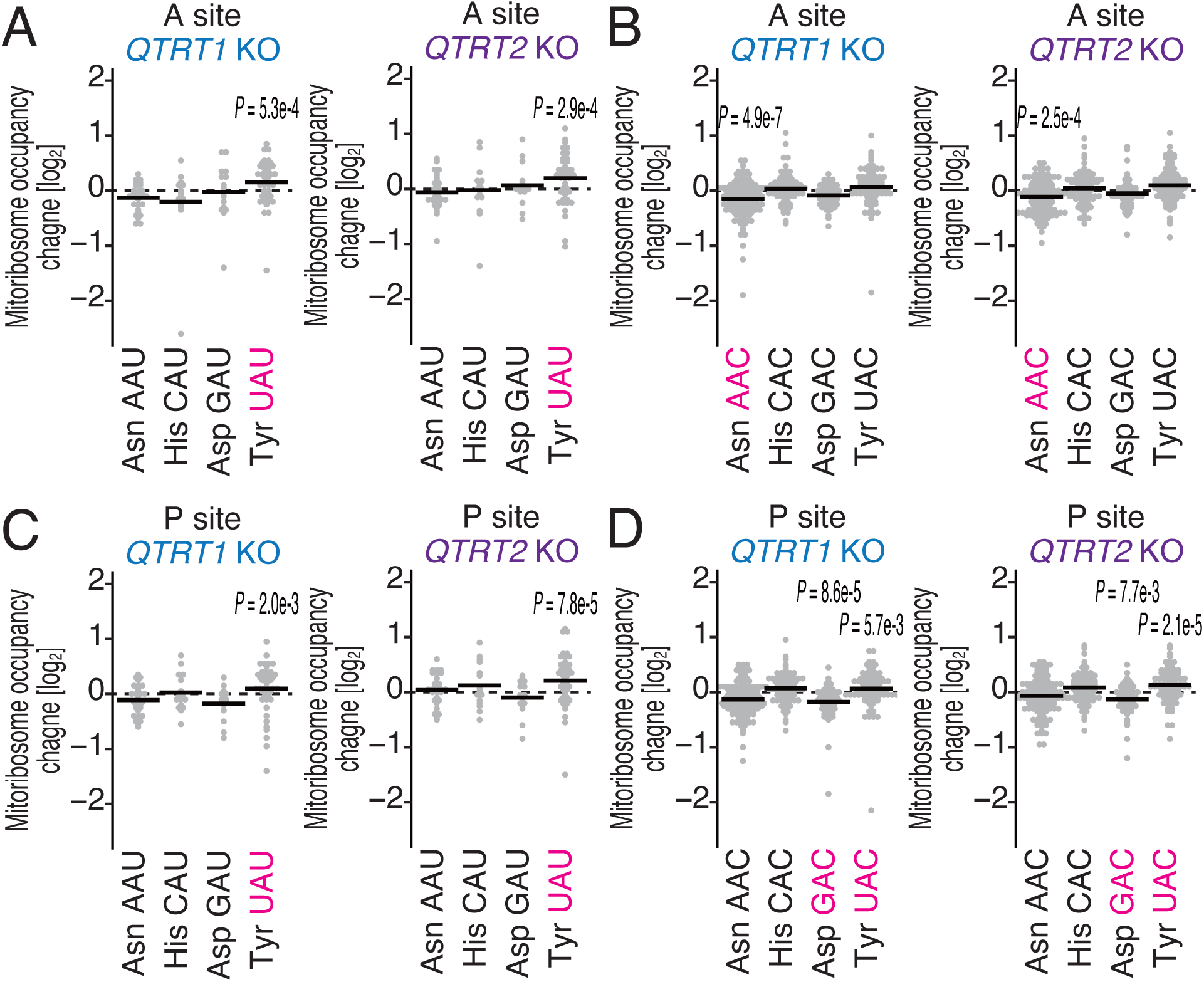
Characterization of MitoIP-Thor-Ribo-Seq data for *QTRT1* KO and *QTRT2* KO cells, related to Figure 4. (A and B) Mitoribosome occupancy changes at A-site U-ending codons (A; AAU, CAU, GAU, and UAU) and C-ending codons (B; AAC, CAC, GAC, and UAC) in *QTRT1* KO and *QTRT2* KO cells. Codons with statistically significant changes in both *QTRT1* KO and *QTRT2* KO are highlighted. (C and D) Mitoribosome occupancy changes at P-site U-ending codons (C; AAU, CAU, GAU, and UAU) and C-ending codons (D; AAC, CAC, GAC, and UAC) in *QTRT1* KO and *QTRT2* KO cells. Codons with statistically significant changes in both *QTRT1* KO and *QTRT2* KO are highlighted. P values were calculated by the Mann‒Whitney *U* test.

**Figure S8.**
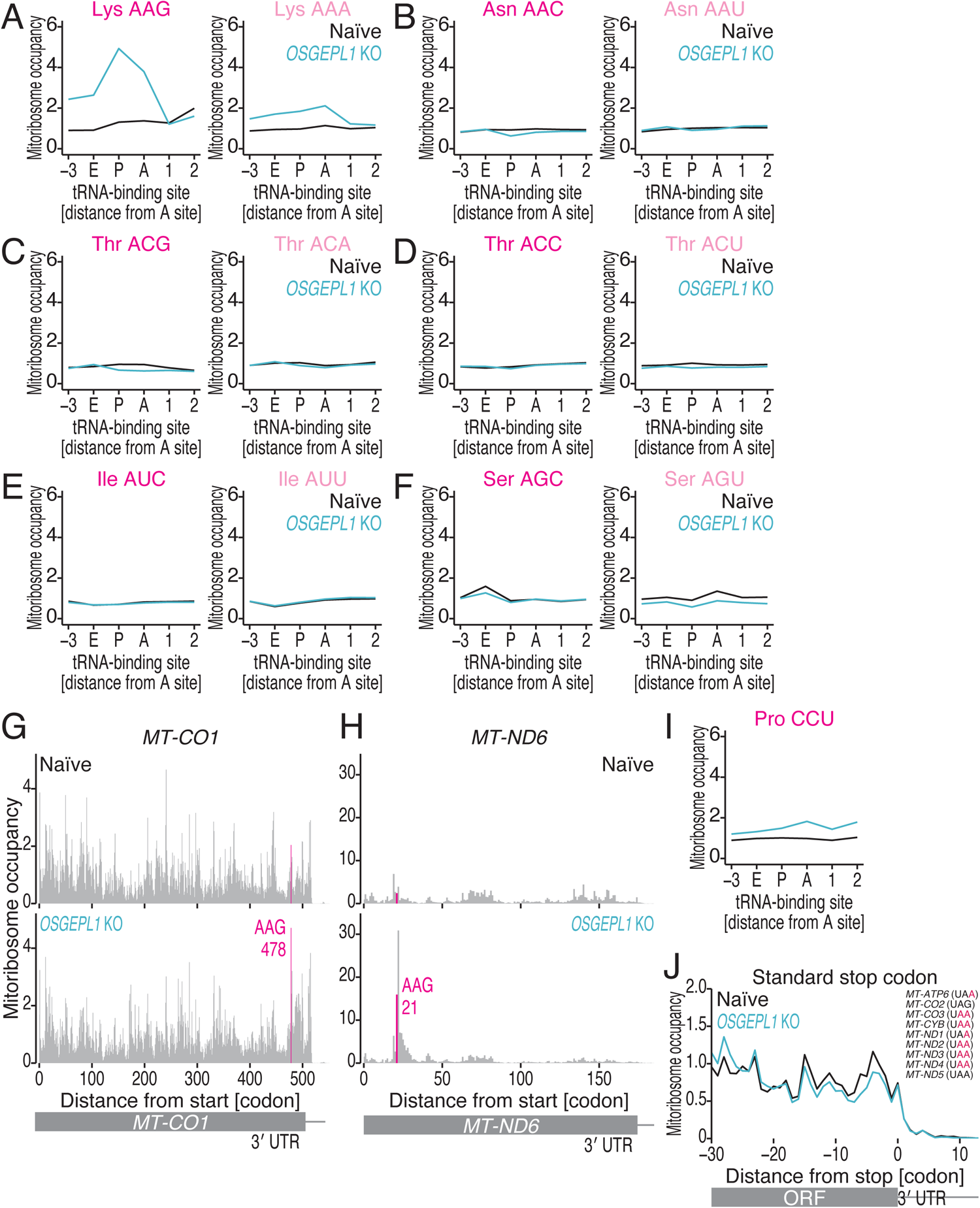
Characterization of MitoIP-Thor-Ribo-Seq data for *OSGEPL1* KO cells, related to Figure 4. (A-F and I) Mitoribosome occupancy of naïve and *OSGEPL1* KO cells for the indicated codons. The scores at each codon position (relative to the A site) are shown. (G and H) Distribution of mitoribosome footprints from naïve and *OSGEPL1* KO cells along the indicated transcripts. The A-site position of each read is indicated. AAG codons are highlighted. (J) Metagene plots for mitoribosome footprints from naïve and *OSGEPL1* KO cells around stop codons. *MT-ATP8* and *MT-ND4L* were excluded from the analysis because their ORF stop codons overlapped with those of *MT-ATP6* and *MT-ND4,* respectively. *MT-CO1* and *MT-ND6* were also excluded from the analysis since they possess noncanonical stop codons (AGA and AGG).

**Figure S9.**
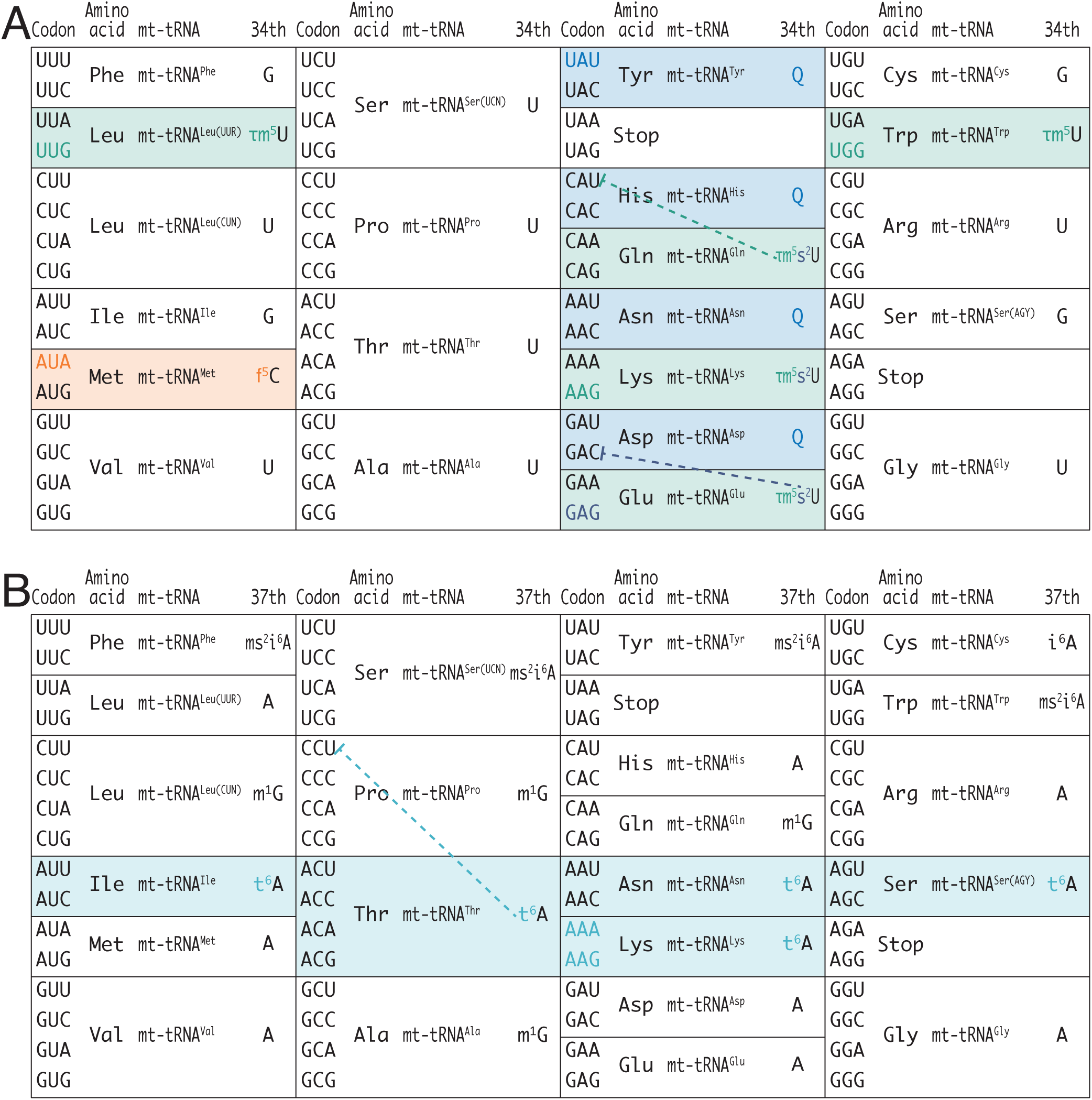
Summary of the impact of mt-tRNA modification on translation elongation in mitochondria, related to Figures 4 and 5. (A) Codon table with the nucleotide of the 34th positions in the corresponding mt-tRNAs. Color-highlighted codons are regulated by the corresponding modifications. Dashed lines indicate that the modifications function to suppress the competition with other mt-tRNA. (B) Codon table with the nucleotide of the 37th positions in the corresponding mt-tRNAs. Color-highlighted codons are regulated by the corresponding modifications. Dashed lines indicate that the modifications function to suppress the competition with other mt-tRNA.

**Figure S10.**
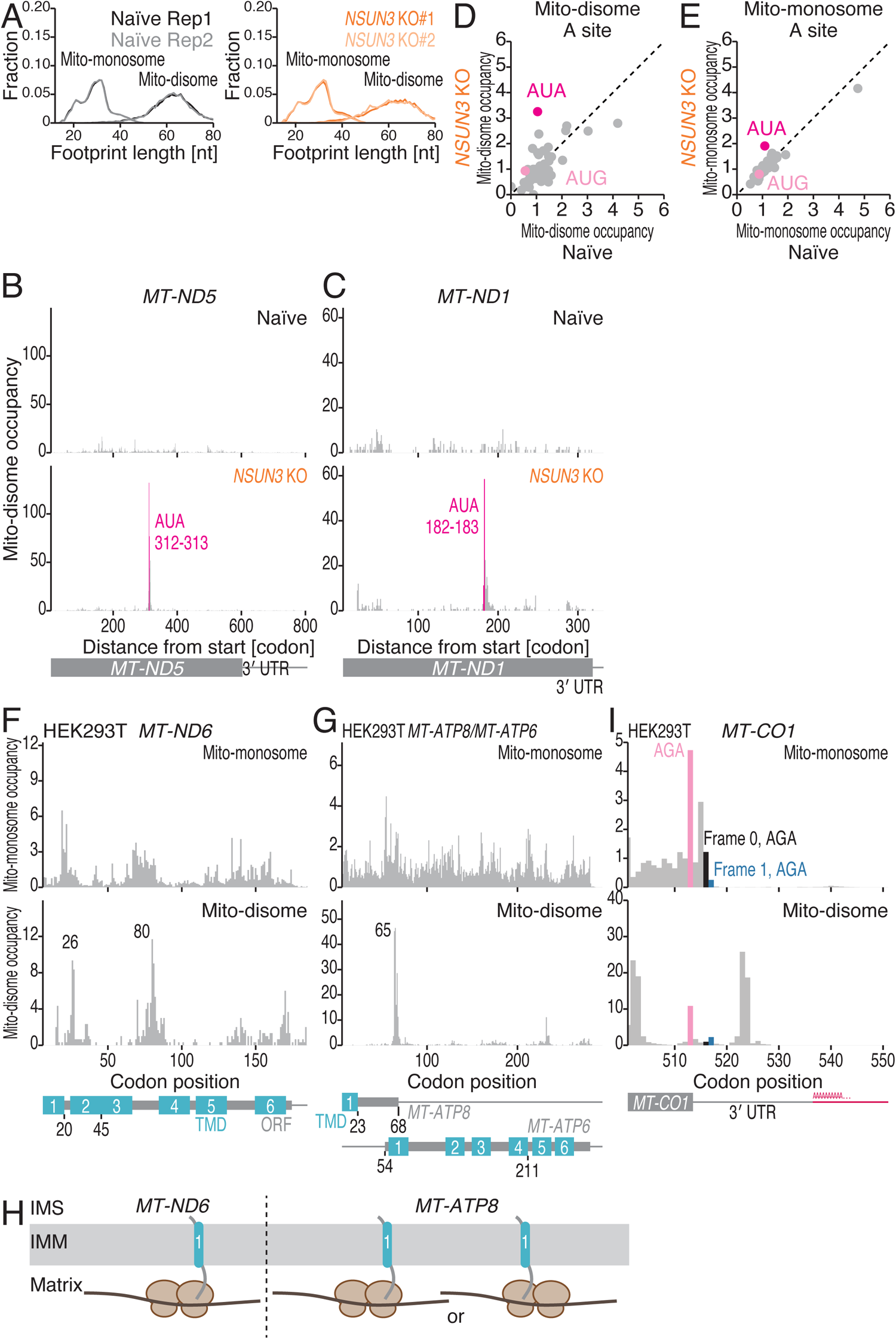
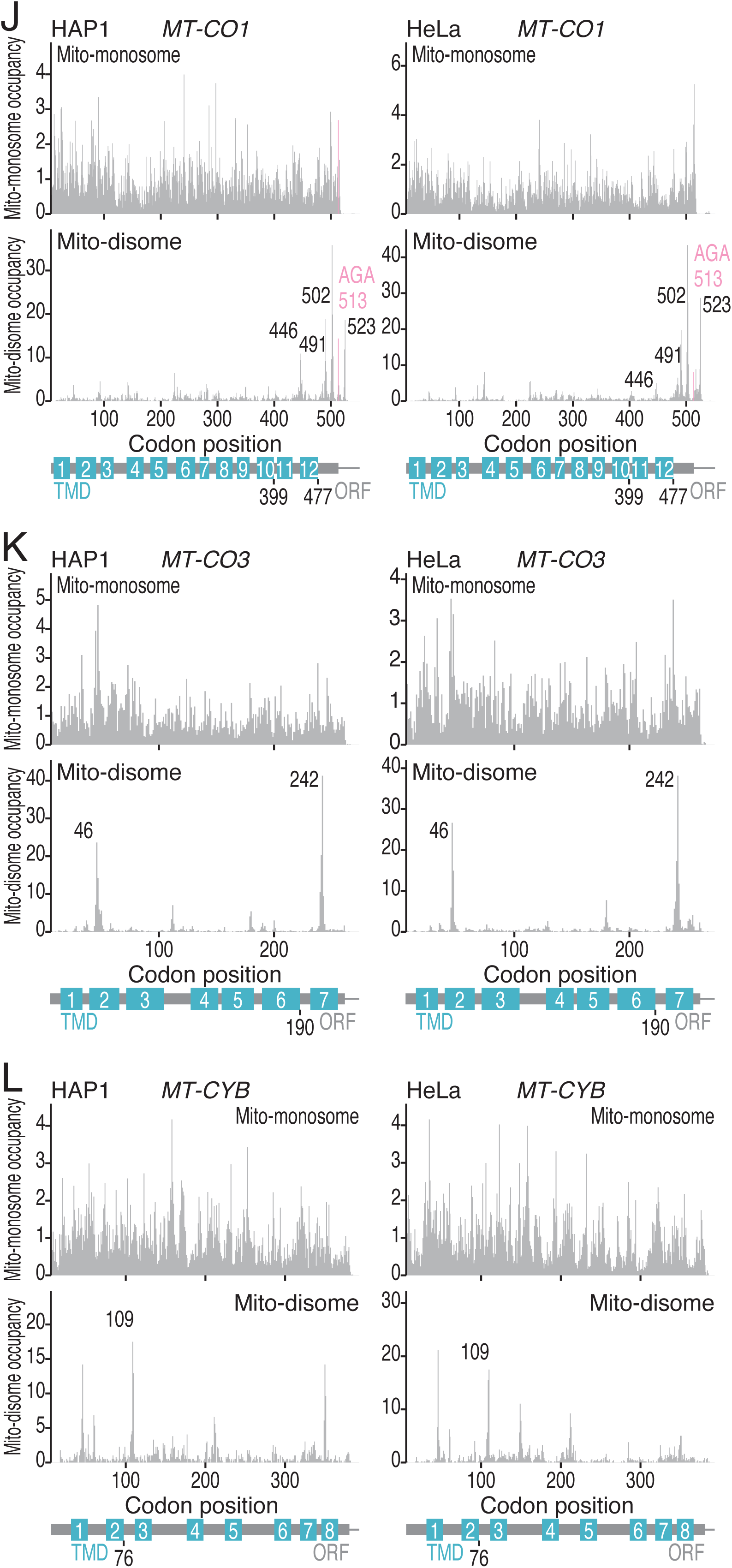
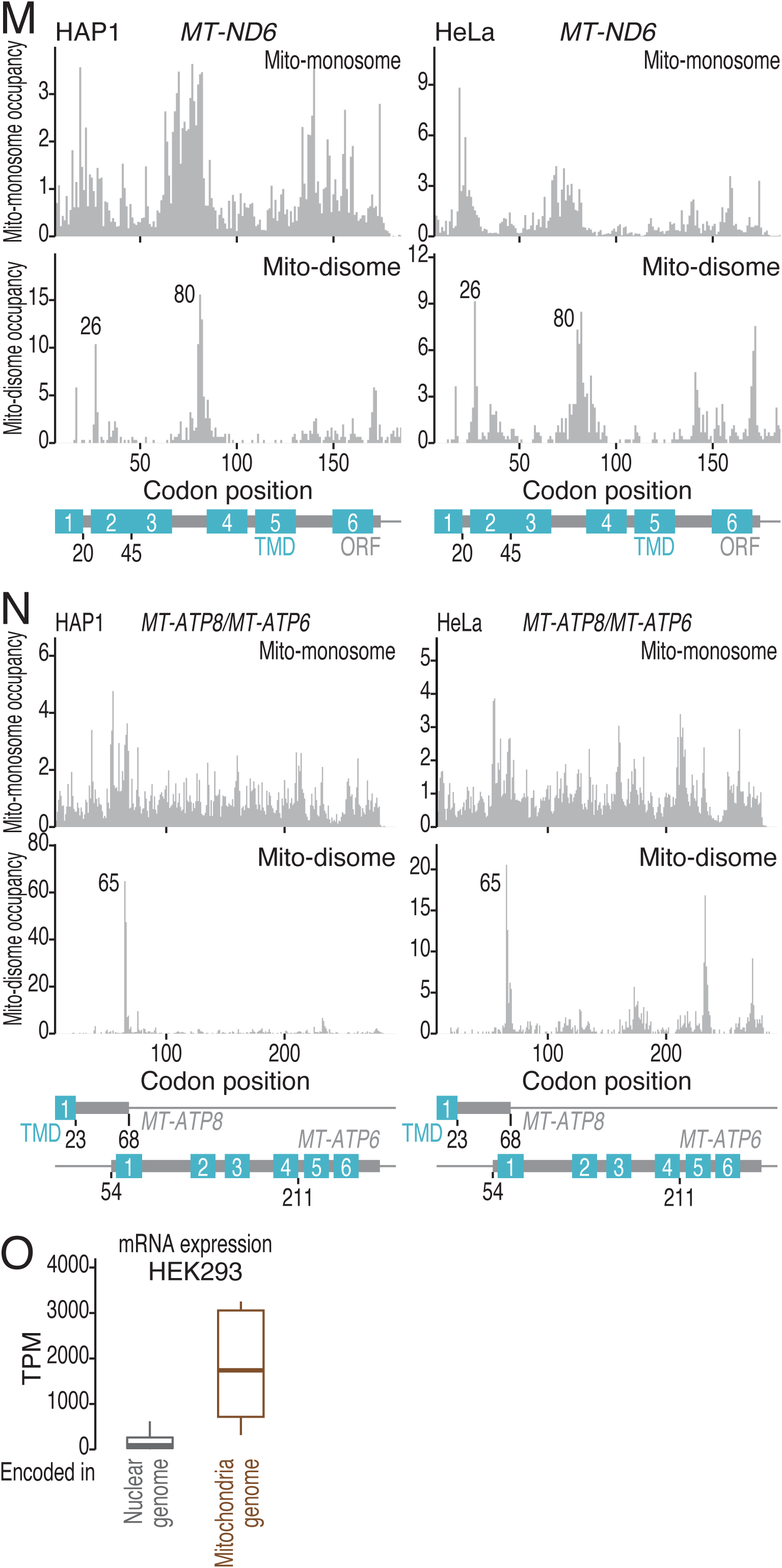
Characterization of MitoIP-Thor-Disome-Seq data, related to Figure 6. (A) The fraction of mito-monosome and mito-disome footprints along the read length for the indicated conditions. The mito-monosome data are the same as those shown in Figure S4G. (B and C) Distribution of mito-disome footprints from naïve and *NSUN3* KO cells along the indicated transcripts. The A-site position (in the case of the mito-disome, the leading paused mitoribosome) of each read is indicated. AUA codons are highlighted. (D and E) Comparison of mito-monosome (E) and mito-disome (D) occupancy at A-site codons under the indicated conditions. (F-G) Distribution of mito-monosome and mito-disome footprints from naïve HEK293 cells along the indicated transcripts. The A-site position (in the case of the mito-disome, the leading paused mitoribosome) of each read is indicated. (H) Schematic of the configurations of the mito-disome pause sites found in *MT-ND6* and *MT-ATP8/ATP6* with cotranslational membrane insertion of their nascent peptides. (I) Zoomed-up view of Figure 6E around the stop codon. The out-of-frame stop codons are highlighted. (J-N) Distribution of mito-monosome and mito-disome footprints from naïve HAP1 cells and HeLa cells along the indicated transcripts. The A-site position (in the case of the mito-disome, the leading paused mitoribosome) of each read is indicated. (O) Box plot of the mRNA expression of nuclear genome-encoded genes and mitochondrial genome-encoded genes in HEK293 cells. The median (centerline), upper/lower quartiles (box limits), and 1.5× interquartile range (whiskers) are shown. TPM, transcripts per million.

**Table S1. Summary of deep sequencing data used in this study, related to Figures 1-6.** For each dataset, the library name, treatment, species name, cell line or strain name, rRNA depletion method, Ribo-Calibration application, replicate number, data type, RNA-Seq kit name, input amount, reference, and accession number are listed.

**Table S2.**
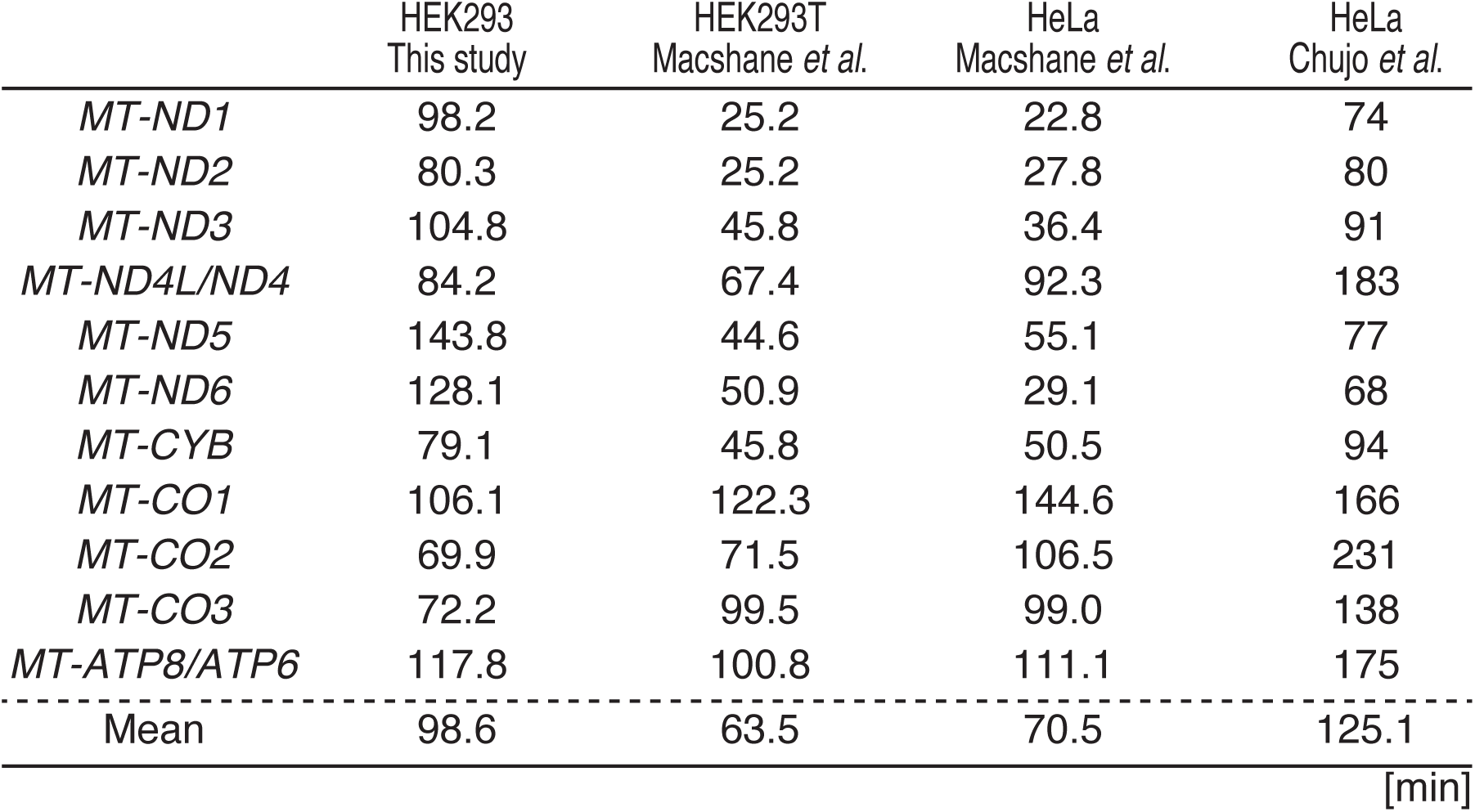
Summary of mRNA half-lives reported by earlier studies and this study, related to Figures 2. mRNA half-lives (min) were measured by BRIC-Seq (this study for HEK293 cells), TimeLapse-seq (for HEK293T and HeLa cells) ^55^, and transcription shut-off by ethidium bromide and chase (for HeLa cells) ^54^.

|  | HEK293<br>This study | HEK293T<br>Macshane *et al.* | HeLa<br>Macshane *et al.* | HeLa<br>Chujo *et al.* |
| --- | --- | --- | --- | --- |
| *MT-ND1* | 98.2 | 25.2 | 22.8 | 74 |
| *MT-ND2* | 80.3 | 25.2 | 27.8 | 80 |
| *MT-ND3* | 104.8 | 45.8 | 36.4 | 91 |
| *MT-ND4L/ND4* | 84.2 | 67.4 | 92.3 | 183 |
| *MT-ND5* | 143.8 | 44.6 | 55.1 | 77 |
| *MT-ND6* | 128.1 | 50.9 | 29.1 | 68 |
| *MT-CYB* | 79.1 | 45.8 | 50.5 | 94 |
| *MT-CO1* | 106.1 | 122.3 | 144.6 | 166 |
| *MT-CO2* | 69.9 | 71.5 | 106.5 | 231 |
| *MT-CO3* | 72.2 | 99.5 | 99.0 | 138 |
| *MT-ATP8/ATP6* | 117.8 | 100.8 | 111.1 | 175 |
| Mean | 98.6 | 63.5 | 70.5 | 125.1 |

[min]

**Table S3.**
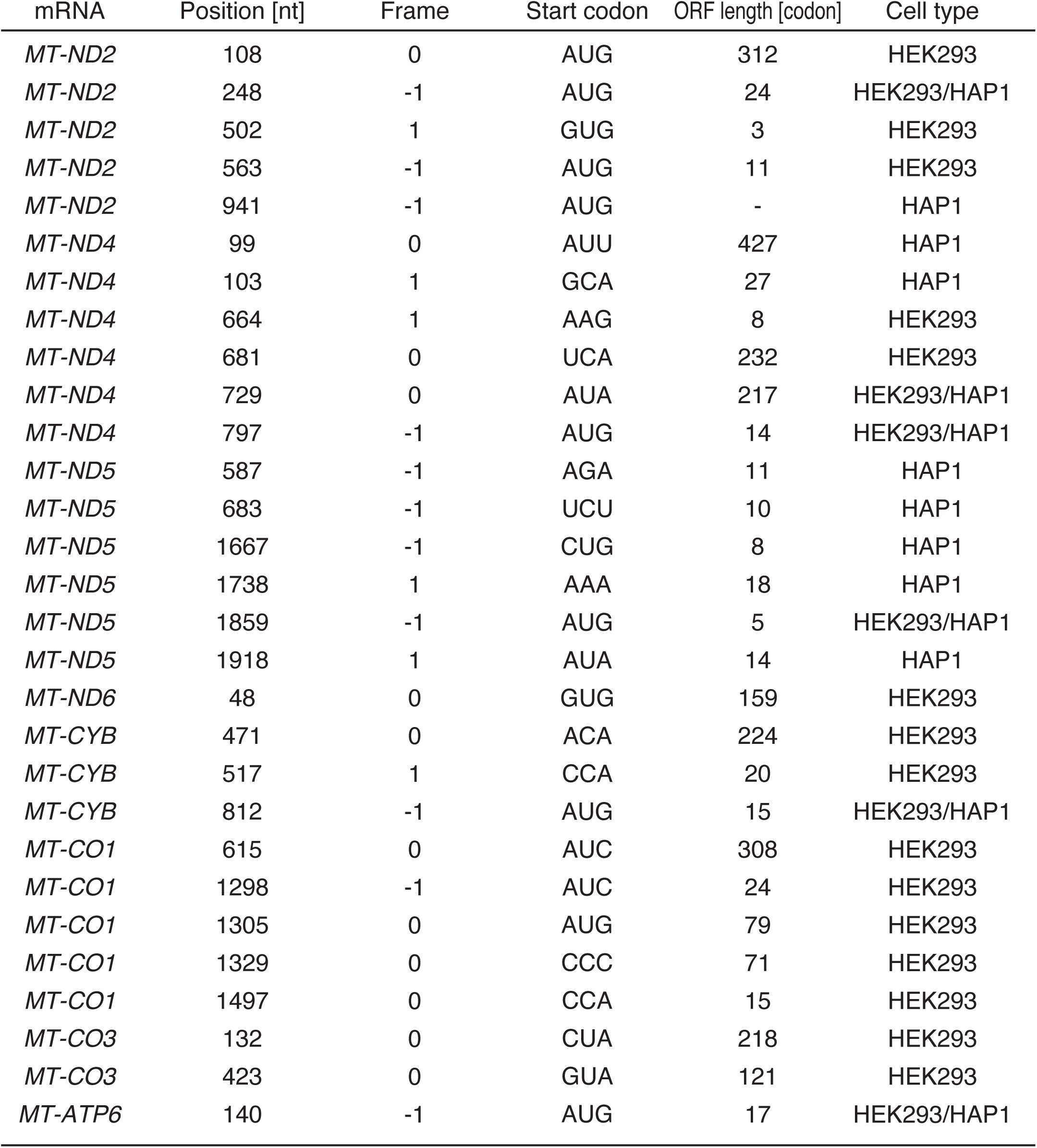
Summary of predicted ORFs translated by cryptic initiation sites, related to Figure 3. Cryptic translation initiation sites are listed with transcripts, the position of the first nucleotide of the cryptic initiation codons from the 5′ ends the transcript (the first nucleotide of the transcript was set to 0), the codon sequences, predicted ORF lengths, and cell-type dependency.

## Notes

### Summary of Updates

Figure 1-6 updated; Figure S1-10 updated; texts revised according to the figure update.

